# Pan-azole- and multi-fungicide-resistant *Aspergillus fumigatus* is widespread in the United States

**DOI:** 10.1101/2023.12.14.571763

**Authors:** BN Celia-Sanchez, B Mangum, LF Gómez Londoño, C Wang, B Shuman, MT Brewer, M Momany

## Abstract

*Aspergillus fumigatus* is an important global fungal pathogen of humans. Azole drugs are among the most effective treatments for *A. fumigatus* infection. Azoles are also widely used in agriculture as fungicides against fungal pathogens of crops. Azole-resistant *A. fumigatus* has been increasing in Europe and Asia for two decades where clinical resistance is thought to be driven by agricultural use of azole fungicides. The most prevalent mechanisms of azole resistance in *A. fumigatus* are tandem repeats (TR) in the *cyp51A* promoter coupled with mutations in the coding region which result in resistance to multiple azole drugs (pan-azole resistance). Azole-resistant *A. fumigatus* has been isolated from patients in the United States (U.S.), but little is known about its environmental distribution. To better understand the distribution of azole-resistant *A. fumigatus* in the U.S., we collected isolates from agricultural sites in 8 states and tested 202 isolates for sensitivity to azoles. We found azole-resistant *A. fumigatus* in agricultural environments in 7 states showing that it is widespread in the U.S. We sequenced environmental isolates representing the range of U.S. sample sites and compared them with publicly available environmental worldwide isolates in phylogenetic, principal component, and ADMIXTURE analyses. We found worldwide isolates fell into three clades and that TR-based pan-azole resistance was largely in a single clade that was strongly associated with resistance to multiple agricultural fungicides. We also found high levels of gene flow with clear recombination between two clades highlighting the potential for azole-resistance to continue spreading in the U.S.

**Importance:** *Aspergillus fumigatus* is a fungal pathogen of humans that causes over 250,000 invasive infections each year. It is found in soils, plant debris and compost. Azoles are the first line of defense antifungal drugs against *A. fumigatus*. Azoles are also used as agricultural fungicides to combat other fungi that attack plants. Azole-resistant *A. fumigatus* has been a problem in Europe and Asia for twenty years and has recently been reported in patients in the U.S. Until this study we didn’t know much about azole-resistant *A. fumigatus* in agricultural settings in the U.S. In this study we isolated azole-resistant *A. fumigatus* from multiple states and compared it to isolates from around the world. We show that *A. fumigatus* that is resistant to azoles and to other strictly agricultural fungicides is widespread in the U.S.

## Introduction

*Aspergillus fumigatus* is an opportunistic fungal pathogen of humans found in soils, plant debris and compost around the world. Causing over 250,000 invasive aspergillosis (IA) cases globally each year in immunocompromised individuals, *A. fumigatus* has a mortality rate approaching 90% with antimicrobial resistance (1, 2). In recognition of its clinical importance the World Health Organization (WHO) has designated *A. fumigatus* one of four critical fungal pathogens (1).

Infections caused by *A. fumigatus* are often treated with azole drugs (3). Azoles are also widely used as agricultural fungicides to combat fungal pathogens of crops (4). Azoles target a conserved Cyp51 protein (also known as Cyp51A, Cyp51B, ERG11) in the pathway that makes ergosterol, a fungal sterol responsible for cell membrane fluidity (5). Binding of azoles to Cyp51 causes arrested growth, toxic sterol intermediates, and fungal cell death (6). *A. fumigatus* has evolved multiple resistance mechanisms to azoles including tandem repeats (TR) in the *cyp51A* promoter to increase expression coupled with point mutations that affect the binding of azoles. Tandem repeats of 34-, 46- and 53-bases in the promoter coupled with point mutations in the coding region of *cyp51A* cause pan-azole resistance, defined as high levels of resistance to multiple azole drugs (7–9). The TR_34_/L98H and TR_46_/Y121F/T289A alleles are most associated with pan-azole resistance (10). Some isolates with TR in the *cyp51A* promoter have also been found to carry alleles for resistance to agricultural fungicides of the quinone outside inhibitor (QoI), benzimidazole (MBC) and succinate dehydrogenase inhibitor (Sdh) classes (11).

Previous studies have shown that worldwide *A. fumigatus* isolates are not structured geographically or by clinical or environmental source of isolation. However, two clades are often identified: Clade A, where isolates with multifungicide resistance and TR mutations causing azole resistance are clustered; and Clade B, which contains all other isolates (11–14). Recently, Lofgren et al. identified a third clade of *A. fumigatus* in a pan-genome analysis and renamed the existing clades with numbers : Clade 1 (Clade B), Clade 2 (Clade A), and Clade 3, which was previously called *A. fumigatus sensu stricto* (15, 16). Clade 3 contains mostly azole sensitive isolates (16, 17).

Azole-resistant *A. fumigatus* has been recognized as a clinical problem in Europe and Asia for two decades (18, 19). Abundant evidence strongly supports the view that agricultural use of azoles led to pan-azole clinical resistance (11, 14, 18), including genetic similarities in azole-resistant *A. fumigatus* isolates from agricultural settings where azoles are used as fungicides and from clinical settings where azoles are used as drugs (14) and the presence in clinical isolates of markers for resistance to fungicides only used in agriculture (11). Resistance to clinical azole drugs has been reported in the U.S. (13) where azoles are also used as agricultural fungicides (4, 10); however, information about the presence of azole-resistant *A. fumigatus* in agricultural environments in the U.S. has been limited. Environmental sampling of a cull pile from a single agricultural site in Georgia yielded the first environmental TR *A. fumigatus* strains identified in the U.S. (20). This was followed by more widespread sampling in Florida and Georgia where 12 of the 123 isolates were found to be pan-azole resistant (11).

Analysis of 179 U.S. isolates by the Centers for Disease Control and Prevention (CDC) including passive surveillance data from clinical settings in 38 states, 13 environmental isolates from a Georgia peanut pile, and two historical isolates from the CDC collection showed that 26% of isolates were resistant to azoles (13). To better understand azole-resistant *A. fumigatus* in the U.S., we collected samples from agricultural settings in previously unsampled states on the east and west coasts, isolated *A. fumigatus*, tested isolates for sensitivity to azoles and for markers of agricultural fungicide resistance and analyzed their genetic relationships with each other and with worldwide environmental and clinical isolates using whole genome sequence data. We found azole-resistant *A. fumigatus* in field soil and plant debris from different locations and different crops cultivated in both the east and west coasts of the U.S. Our population genomic analyses support the existence of 3 clades with a strong association of TR-based azole resistance and multi-fungicide resistance with Clade 2. Moreover, we show recombination between Clades 1 and 2, which suggests that pan-azole resistance will keep spreading within U.S. populations of *A. fumigatus* as it has worldwide.

## Methods

### Sampling

Sampling was conducted as previously described (11). Briefly, between November 2018 and October 2019 soil, plant debris, or compost was collected from 29 agricultural sites in the eastern U.S. (New York, Pennsylvania, Virginia, South Carolina, and Georgia), and from 23 agricultural sites in the western U.S. (Washington, Oregon, and California) (Table 1). Soil and compost were sampled by taking 3-5 soil cores to a depth of 10-15 cm. Fallen plant debris was collected with the soil if present. Collections were limited to 4 samples per site to minimize isolation of clones. Samples were shipped over-night in sealed plastic bags, and then stored at 4°C and with bags open to allow for gas exchange.

**Table 1:**
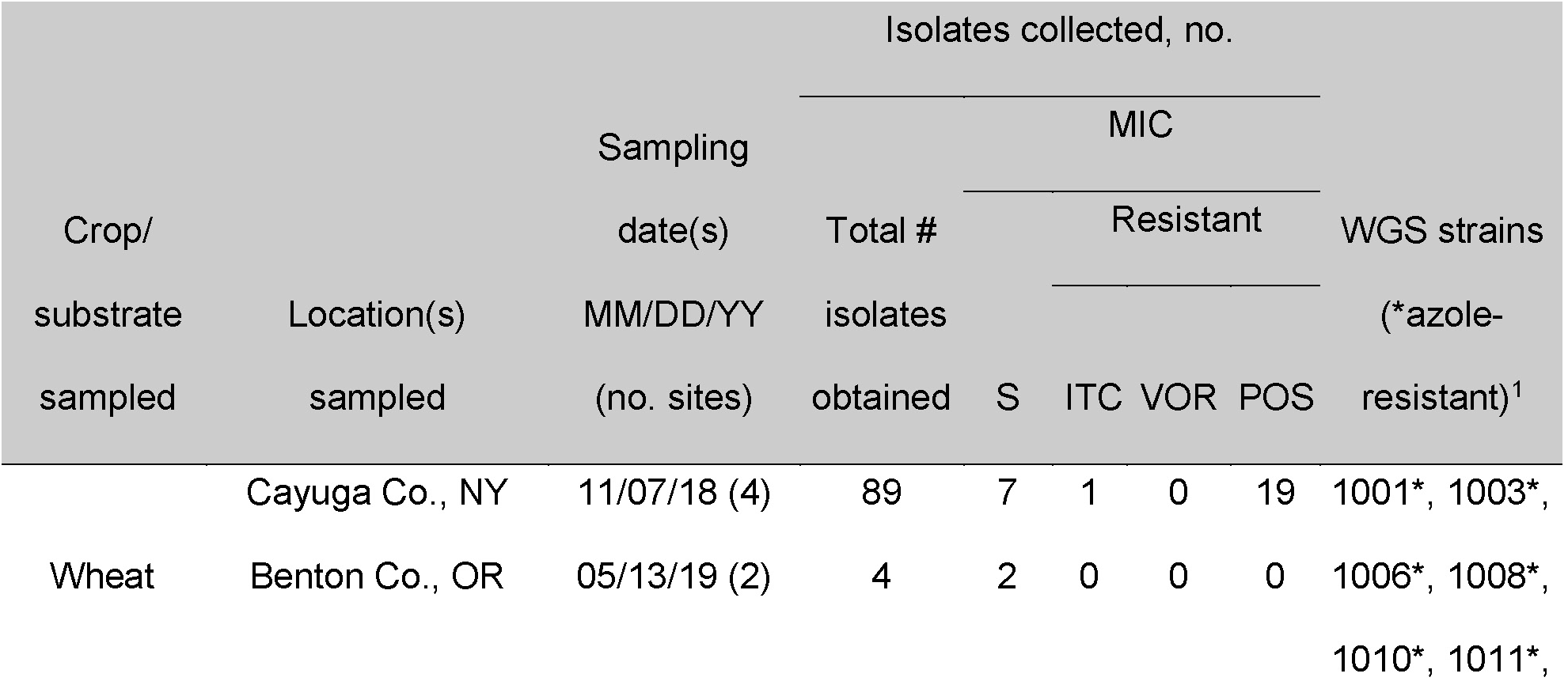
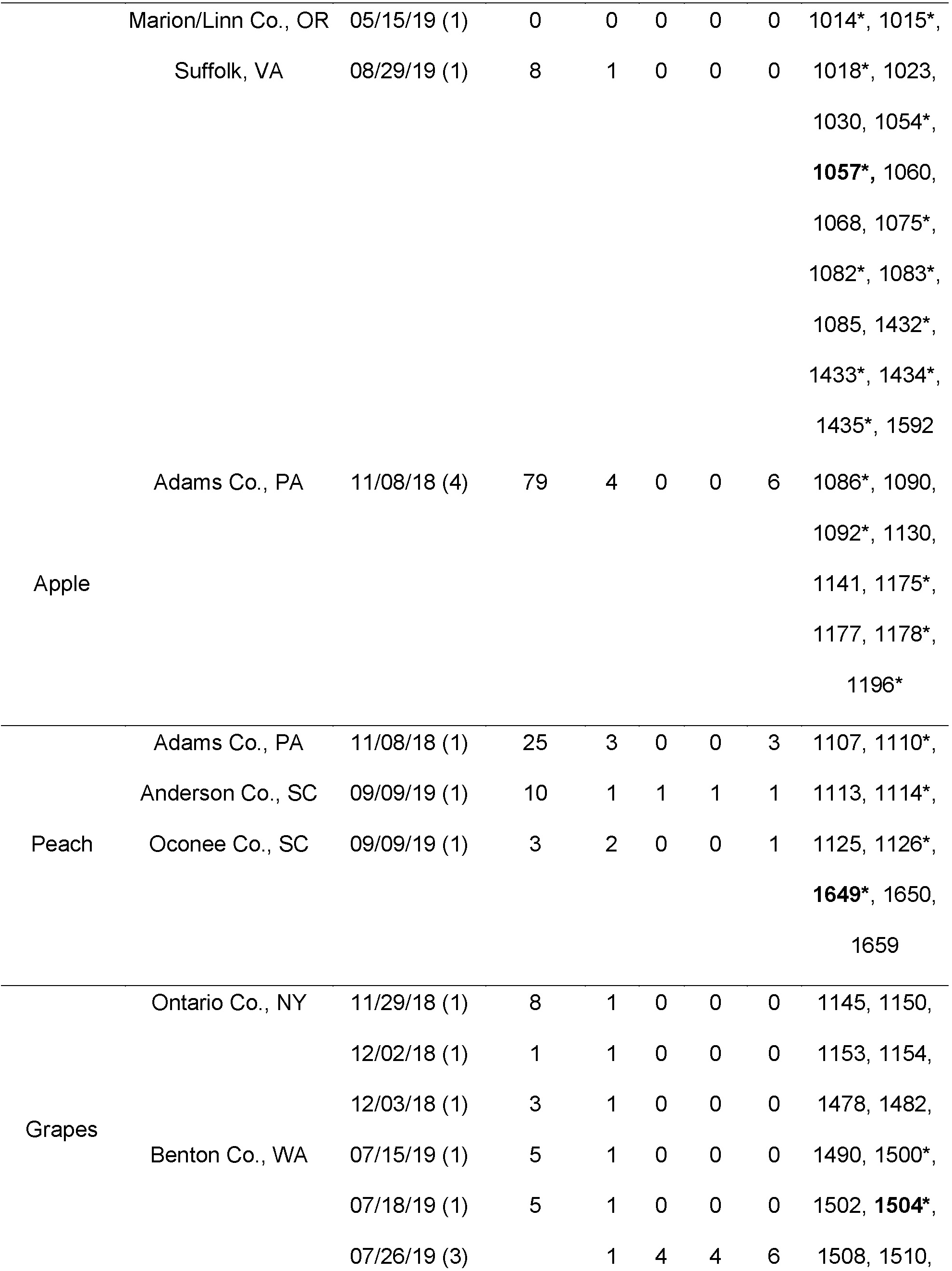
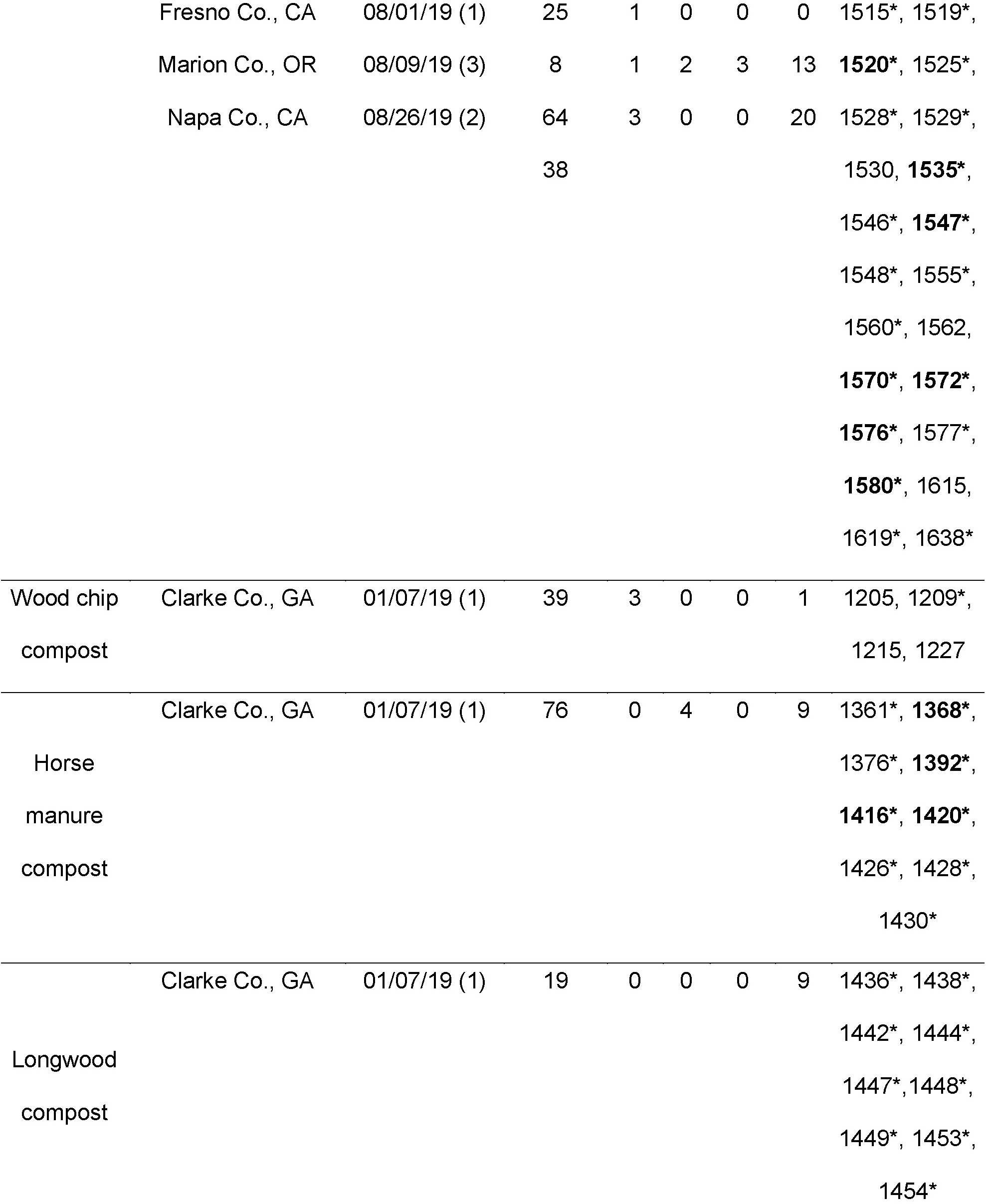
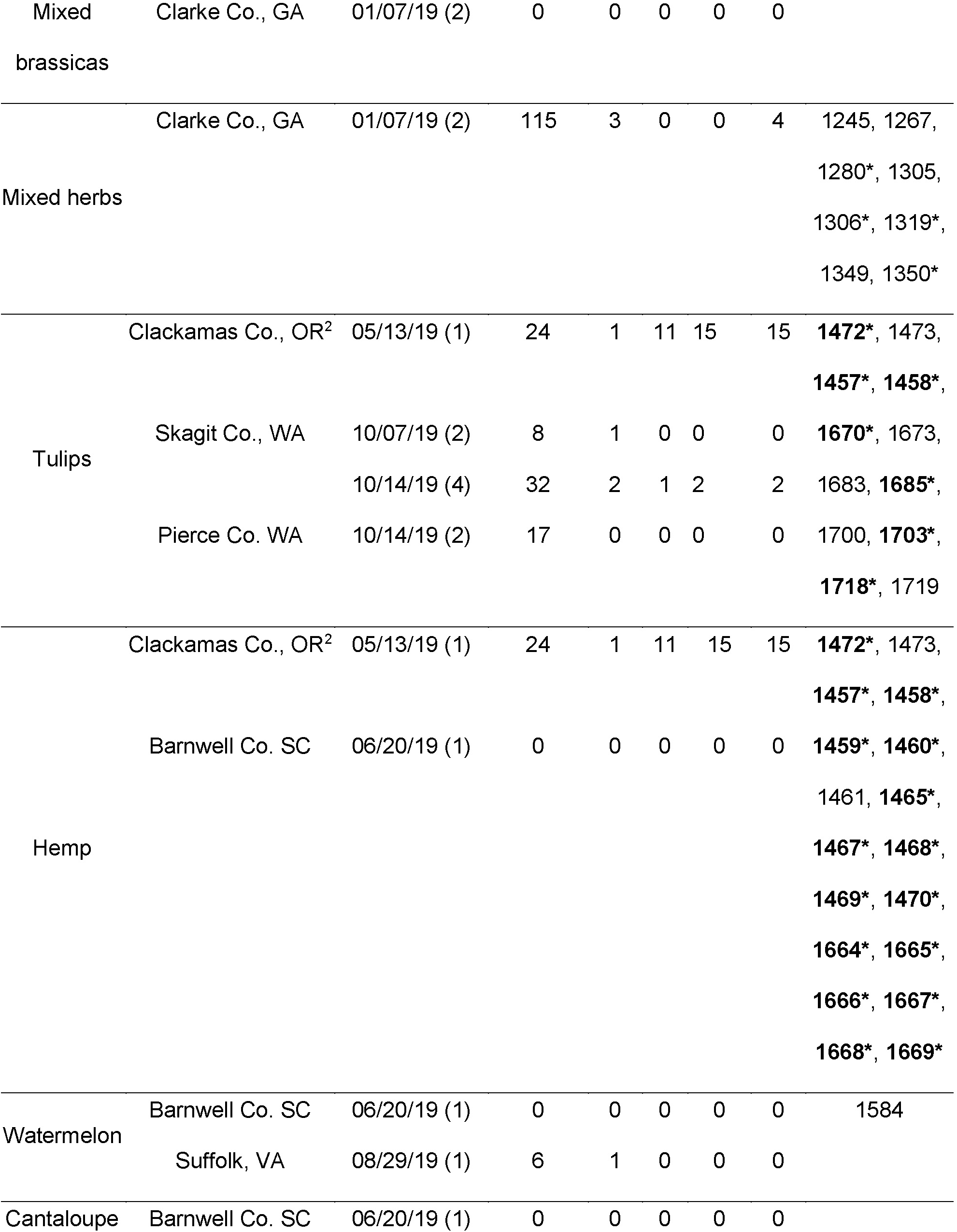
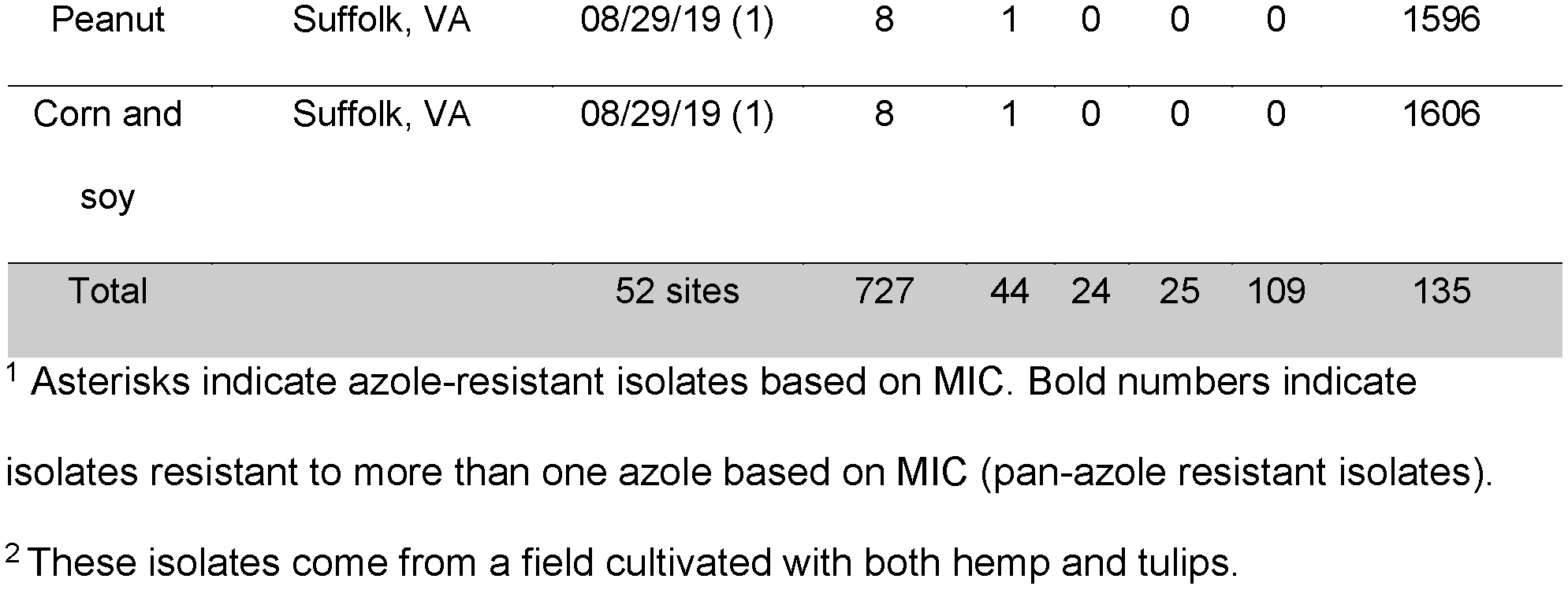
Sampling of *A. fumigatus* strains from agricultural sites.

### Isolation and storage

The samples were processed as described previously with some modifications (11, 18, 20). Briefly, 2 g of soil was suspended in 8 ml of 0.1 M sterile sodium pyrophosphate. Samples were vortexed for 30 sec and allowed to settle for 1 minute. From the supernatant, 100 µl was pipetted onto 10 plates each of Sabouraud dextrose agar (SDA) control plates or SDA supplemented with 3 µg/mL of the fungicide tebuconazole or with 4 µg/mL of the antifungal itraconazole. All plates additionally contained 50 µg/mL of chloramphenicol and 5 µg/mL of gentamicin to inhibit bacterial growth, and 25 µg/mL of Rose-Bengal dye to reduce the growth of *Rhizopus* and *Mucor*. The plates were sealed with micropore tape and incubated at 45°C for 48-72 hours. Colonies of *A. fumigatus* were identified by morphology. Colonies that grew on azole-drug amended media were single colony isolation streaked on fresh azole-containing media without other antimicrobials added. Several colonies that grew on negative control plates were single colony isolation streaked on azole-drug amended media to identify sensitive isolates for comparison in whole genome sequencing (WGS) analyses. All collected isolates were stored in 15% glycerol at -80°C.

### Minimum inhibitory concentration (MIC) assays for antifungal susceptibility testing

Using the Clinical Laboratory Standard Institute broth microdilution method for antifungal susceptibility testing, we tested 160 environmental *A. fumigatus* isolates for sensitivity to the fungicide tebuconazole (TEB; TCI America, Oregon, USA), and the antifungals itraconazole (ITC; Thermo Sci Acros Organics, New Jersey, USA), voriconazole (VOR; Thermo Sci Acros Organics, New Jersey, USA), and posaconazole (POS; Apexbio Technology, Texas, USA). The isolates were grown on complete media slants for 4 days and were then harvested using 3 mL of 0.05% Tween-20 to suspend the hydrophobic spores in solution. The spore suspensions were adjusted to an optical density of 0.09 – 0.13 at 530 nm using a spectrophotometer and, 100 µL of spore suspension were added to a microtiter plate well containing100 ul of RPMI 1640 liquid medium (Thermo Sci Gibco, California, USA) and azoles with final concentrations ranging from 0 to 16 μg/ml. The plates were incubated for 48 hours at 37°C, and then characterized by the European Committee on Antibiotic Susceptibility Testing (EUCAST) breakpoints (21). Minimum inhibitory concentration (21) break points were defined as the lowest concentration where 100% of growth was inhibited. The assay was performed twice on all 160 isolates. For instances where MIC break points differed by greater than one dilution between replicates, the isolates were assayed a third time. For classification of sensitivity or resistance for TEB (>3 µg/mL), ITC (> 2 µg/mL), VOR (> 2 µg/mL), and POS (> 0.25 µg/ml, if = to 0.25µg/mL consider resistant if ITC is resistant), we used the current recommended clinical breakpoints of antifungal resistance for *A. fumigatus* published by EUCAST 2020 (21, 22).

### DNA Extraction and Sequencing

Genomic DNA was extracted from *A. fumigatus* isolates using a CTAB protocol described previously with some modifications (11, 23). Briefly, cultures were grown overnight in complete medium broth and approximately 200 mg of mycelium was collected and transferred to 2 ml microcentrifuge tubes containing 150 mg of glass disruption beads and three 3-mm steel beads and lyophilized for 24 hours. Lyophilized cells were disrupted using GenoGrinder 2010 (OPS Diagnostics, Lebanon, NJ) at 1750 rpm for 30 seconds. To the pulverized tissue, 1 mL of CTAB lysis buffer (100 mM Tris pH 8.0, 10 mM EDTA, 1% CTAB, 1% BME) was added, and the samples were incubated for 30 minutes at 65°C. After incubation, 300 µl of 5M KAc was added to the samples and incubated on ice for 30 minutes. The supernatant was extracted twice with chloroform, and the aqueous, clear top layer containing DNA was precipitated by addition of an equal volume of ice-cold 100% isopropanol followed by centrifugation.

The DNA pellet was washed in 70% EtOH, then 100% EtOH, and air-dried. The resuspended sample was treated with RNase A according to the JGI RNase A DNA cleanup protocol. DNA was checked for quality and quantified using NanoDrop One (Thermo Sci, New Jersey, USA). Library preparation and Illumina NextSeq 2000 P3 sequencing was conducted at the Georgia Genomics and Bioinformatics Core at the University of Georgia, Athens, GA. Sequences were deposited in NCBI under project PRJNA991533.

### Gene analysis

Whole-genome sequences were assembled for each isolate using SPAdes v3.14.1 with options “–careful” and “–trusted-contigs” (24). Nucleotide BLAST databases were generated for all sequences from SPAdes contig.fasta output using BLAST+ v2.11.0 (25). To investigate genes involved in fungicide resistance, databases were searched by blastn for *A. fumigatus cyp51A* (Afu4g06890), *benA* (Afu1g10910), *cytB* (AfuMt00001), and *sdhB* (Afu5g10370). Blast hits were extracted using BEDtools v2.30.0 (26). Gene analysis was performed using Geneious v2021.0.4.

Genes *MAT1-1-1* (AY898660.1) and *MAT1-2-1* (AFUA_3G06170) indicative of the *MAT1-1* or *MAT1-2* idiomorphs of the mating-type locus (*MAT1*) were used in separate BLAST searches to identify the mating type of each isolate.

To test if Clade 3 was a cryptic species, we used eight housekeeping genes (*act1* (AFUA_6g04740); *efg1* (AFUA_2g02870); *gapdh* (AFUA_5g01970); *hishH4* (AFUA_2g13860); ITS region (AFUA_4g02100); *rpb2* (AFUA_7g01920); *sdh1* (AFUA_3g07810); *tef1* (AFUA_1g06390)) that were identified using a BLAST. As described above, genes were identified and extracted, then analyzed in Geneious v2021.0.4 and aligned to the reference using the “Map to Reference” option with the Geneious mapper, high sensitivity/medium sensitivity, and fine-tuning iterations up to 5 times options. Alignments were used to construct the neighbor-joining gene trees described below.

### Variant calling and phylogenetic analysis

Raw reads were mapped to Af293 (GCF_000002655.1) using BWA v0.7.17 (27). Text pileup outputs were generated for each sequence using SAMtools v1.6 mpileup with option –I to exclude insertions and deletions (28). BCFtools v1.6 call was used to call single nucleotide polymorphisms (SNPs) with options -c to use the original calling method and --ploidy 1 for haploid data (29). Consensus genome sequences in fasta format were extracted from vcf files using seqtk v1.2 (available at https://github.com/lh3/seqtk) with bases with a phred quality score below 40 counted as missing data (N). Because insertions and deletions were removed when mapping reads to the reference, all genome sequences are already mapped to the same coordinates as the reference sequence and there was therefore no need for a multiple alignment step.

MEGA X (30) was used to create a neighbor-joining tree using the Tamura-Nei model with 100 bootstraps for all whole genome sequences. Neighbor joining is a suitable method for constructing whole genome trees from SNP data within a frequently recombining species like *A. fumigatus*. Visualization was done using iTOL (31). MEGA X (30) was used to create neighbor-joining gene trees using the Kimura 2-parameter model (32) with 100 bootstraps for all housekeeping genes.

### Population genetic analyses

Principal-component analysis (PCA) was performed using the smartpca program of EIGENSOFT v7.2.1 (33). PCA results were plotted using the ggplot2 package in R version 4.2.1 (34, 35).All vcf files were merged using BCFtools v1.9 and thinned using VCFtools v0.1.16 with options –thin to have 1 SNP for every 50 nucleotides, --minQ 40 to filter out low quality SNPs, and –plink to output plink files (ped and map). Plink v1.9b was used with the ped and map files as input and option --make-bed to output a bed file to be used in ADMIXTURE (36). To identify ancestry and admixed strains, ADMIXTURE v 1.3 was used in five independent runs (1, 12345, 33333, 54321, and 98765) with 2 to 10 clusters (K) (37). Resulting Q files were used in the Distruct for many K’s feature of CLUMPAK to align cluster labels across different models (38). CLUMPAK results were added to R version 3.5.0 to be plotted (34). The neighbor-joining tree, PCA, critical values (BIC), and loglikelihood values of each K were used to identify the most likely number of clusters (K).

Using ADMIXTURE plot K19 (Figure S3), isolates showing no recombination within subclusters of Clade 1 (Af293-Bowyer, AFIS2275, DI 16-5, eAF1150, eAF1490, F17764) and Clade 2 (C30, C114, C151, C188, eAF234, eAF1500) were used to paint chromosomes of recombinant strains (B11927, eAF1001, eAF1570, eAF1666, L-2-11-4, NRZ-2017-362) with faChrompaint.pl (39). Each recombinant strain was run with six isolates from Clade 1 and six isolates from Clade 2 to identify *in silico* admixture between populations. Resulting nearest.tsv files were used to find the 90th percentile value (0.00136986) to rerun the analyses with the “-M” option.

## Results

To better understand the prevalence of azole-resistant *A. fumigatus* in U.S. agricultural environments, we collected soil, plant debris and compost from 29 agricultural sites in five eastern states (New York, Pennsylvania, Virginia, South Carolina, and Georgia), and from 23 agricultural sites in three western states (Washington, Oregon, and California). Crops cultivated on our sites included grains, fruits, nuts, and ornamentals (Table 1). With the exception of Georgia, no surveys to detect azole-resistant *A. fumigatus* in agricultural settings have been reported for these eight states. We plated samples onto media containing either no azole, the agricultural fungicide TEB or the clinical antifungal drug ITC and isolated a total of 727 isolates of *A. fumigatus*. Following this preliminary selection, we determined the MIC for the clinical antifungal agents ITC, VOR, and POS by broth dilution assays for 202 isolates representing all sites and including azole-sensitive and resistant isolates. Of these isolates, 44 were sensitive to all three azoles tested, 24 were resistant to ITC, 25 were resistant to VOR, 109 were resistant to POS, and 34 were resistant to more than one clinical azole drug (pan-azole resistant) based on EUCAST 2020 cutoffs (Table 1) (21, 22). We performed whole genome sequencing (WGS) on 135 isolates representing the range of collection sites and azole sensitivities.

Previous studies reported on clinical *A. fumigatus* isolated from health care settings in 37 states (13) and environmental *A. fumigatus* isolated from agricultural settings in Florida and Georgia (11, 20). We placed our new isolates and data for 295 publicly available isolates on a map allowing visualization of the distribution of all clinical and environmental *A. fumigatus* so far reported in the U.S. (Figure 1, Table S1). It is impossible to determine the actual frequency of azole-resistant *A. fumigatus* in individual states or associated with particular crops or substrates because sampling intensity and isolate screening and selection has varied widely; however, it is clear that azole-resistant *A. fumigatus* is widespread geographically and across crops and substrates. Pan-azole-resistant isolates were prevalent among samples from grape, compost, tulips and hemp (Table 1). *A. fumigatus* isolates have been reported from 38 states. Of these isolates, 241 were collected from environmental settings in 9 states and 189 from clinical settings in 37 states; 263 were sensitive to clinical azole drugs, 101 to a single azole, and 66 to more than one azole (pan-azole resistant). Including the present study, agricultural sites in only 9 states have been sampled; azole resistant *A. fumigatus* was not detected in agricultural samples from Virginia but was detected in agricultural samples from the other 8 states surveyed (Figure 1, Table S1).

**Figure 1.**
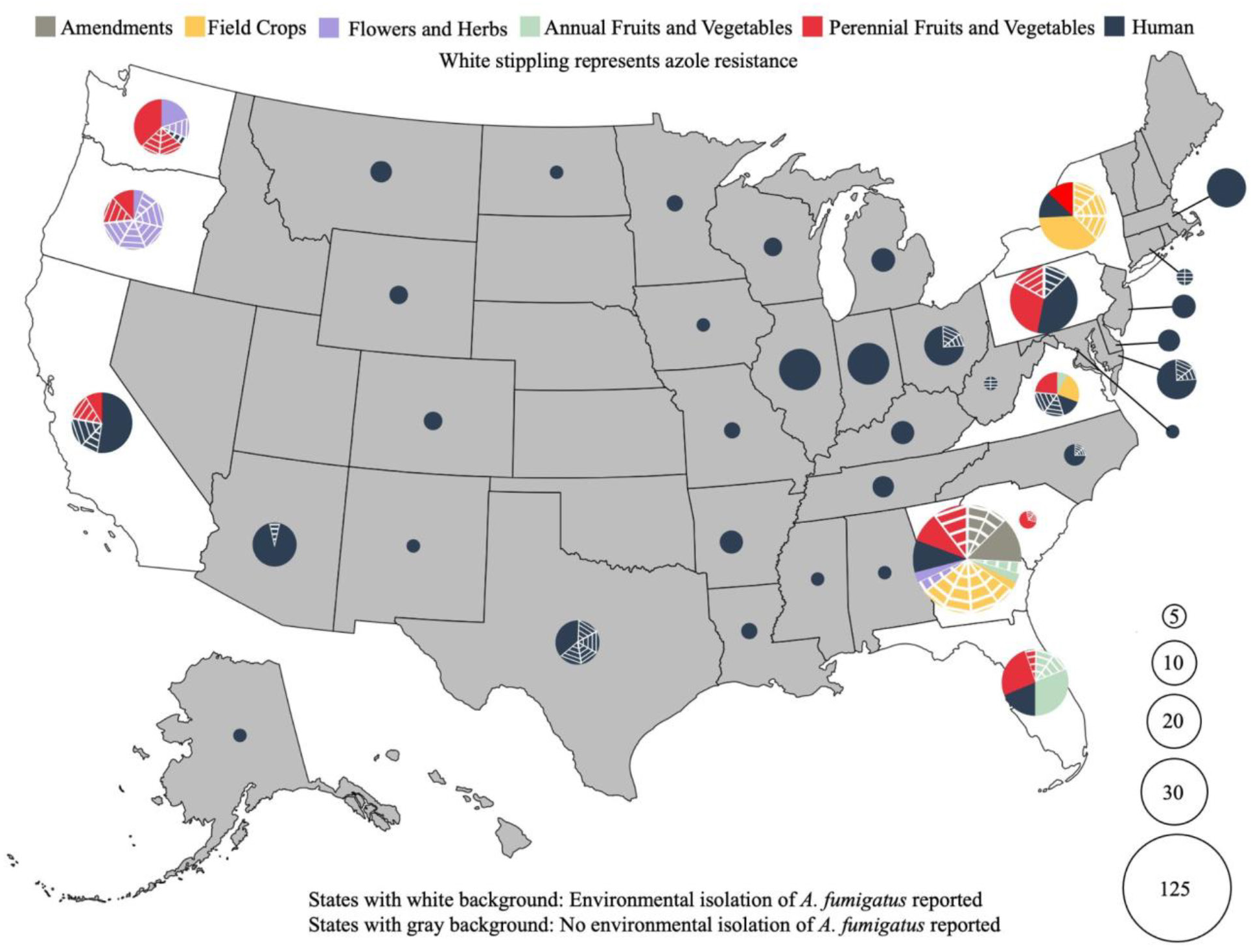
Locations of all *A. fumigatus* isolated from clinical and environmental settings in the U.S. as of June 2023. States with gray background have no reported environmental sampling of *A. fumigatus*. The size of pie charts represents the number of isolates reported in each state. Sampling intensity varied widely, so isolate number does not accurately reflect relative abundance of *A. fumigatus* in each state. Pie chart colors represent sample types (gray: soil amendments; yellow: field crops; purple: flowers and herbs; green: annual fruits and vegetables; red: perennial fruits and vegetables; and dark blue: human clinical). White stripes on pie charts denote azole-resistant isolates. It should be noted that samples in the field crop category in Georgia are primarily from a single peanut cull pile in which multiple azole-resistant isolates were detected. Based on the current study and publicly available data from previous studies as shown in Table S1 (11, 13, 14, 20, 40–48).

Several studies showed that worldwide *A. fumigatus* populations fell into two clusters (Clade A and Clade B) with most TR_34_/L98H and TR_46_/Y121F/T289A-based pan-azole resistance in Clade A (12–14). A recent pan-genome study including more isolates from around the world showed that *A. fumigatus* populations fell into three clusters: Clade 1 contains fewer azole-resistant strains, and appears to be the same as the previously identified Clade B; Clade 2 contains many TR-based pan-azole isolates and appears to be the same as Clade A. The small Clade 3 has no azole-resistance and is made up of 14 clinical isolates from Europe, the U.S. and Canada, and a single environmental isolate from Peru (16). To determine the relationships of U.S. agricultural isolates from the current study to worldwide *A. fumigatus* clinical and agricultural populations, we performed whole genome sequencing on 135 and used the resulting sequences, along with 594 publicly available whole genome sequences to construct a neighbor-joining tree of 729 total sequences (Figure S1, Table S2). To better visualize the tree, 482 representative isolates were chosen for subsampling (Figure 2, Table S2). In addition to analyzing azole-resistance mutations in *cyp51A*, we analyzed genes responsible for resistance to agricultural fungicides (*benA* for MBC class*, cytB* for QoIs, and *sdhB* for SDHI class) and mating-type (*MAT1-1-1*, *MAT1-2-1;* AY898660.1, AFUA_3G06170).

**Figure 2:**
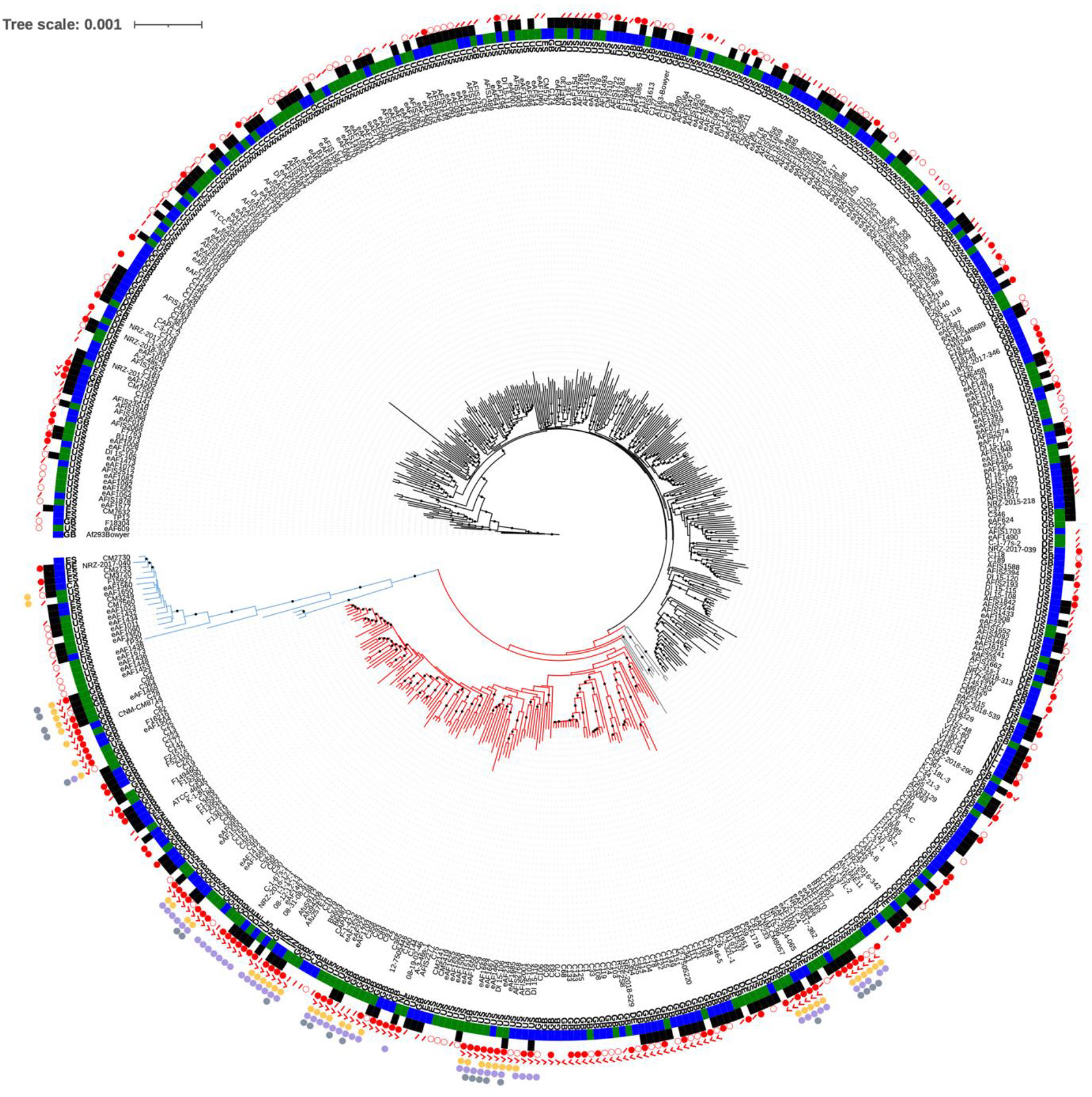
Neighbor-joining tree of 480 subsampled environmental and clinical isolates of *Aspergillus fumigatus.* Whole genome sequences from agricultural sites on the east and west coasts of the United States (eAF1XXX) were analyzed along with publicly available data (Supplementary Table 1). Af293 was used as the reference genome and root of the tree. Bootstrap values greater than 95% are indicated by a dot on the branch. Clade 1 branches are black; Clade 2 branches are red; Clade 3 branches are blue; and branches for strains recombining between Clades 1 and 2 are gray. Country of origin is listed next to each isolate according to their two-letter designation (CA, Canada; DE, Germany; ES, Spain; GB, Great Britain; IN, India; NL, Netherlands; US, United States). Green and blue bars indicate environmental and clinical isolates, respectively. Black and white bars represent mating type *MAT1-1* and *MAT1-2*, respectively. Open red circles indicate azole resistant isolates. Solid red circles indicate pan-azole resistant isolates (i.e., isolates resistant to at least two different clinical azoles based on MIC testing). Red slash marks represent isolates without MIC test data for either itraconazole, voriconazole, or posaconazole. Red check marks indicate isolates with TR mutations. Yellow circles indicate the *cytB* G143A mutation conferring resistance to QoI fungicides (49). Purple circles indicate the *benA* F219Y mutation conferring resistance to MBC fungicides (50). Gray circles indicate the *sdhB* H270Y/R mutations conferring resistance to SDHI fungicides (Fraaije et al. 2012).

Our phylogenetic analysis of 729 worldwide isolates was consistent with three clades with the branch leading to the small Clade 3 showing 100% bootstrap support (Figure 2, Figure S1). *MAT1-1* and *MAT1-2* isolates were present in almost equal proportions throughout the tree (354/729 and 375/729, respectively) and there was no association with clade designation (Figure 2, Figure S1). Based on sequence data and single mutations known to be associated with fungicide resistance phenotypes in *A. fumigatus* (11, 51, 52), we classified isolates as sensitive or resistant to the agricultural fungicides MBC, QoI, and SDHI (Table S2). We used MIC data for ITR, VOR, and POS to determine if isolates were sensitive, resistant to a single clinical azole (non-pan-azole resistant) or to multiple clinical azoles (pan-azole resistant). Because MICs for all three clinical azoles were not reported for all publicly available strains, the actual number of pan-azole-resistant isolates might be higher than our data suggest. Clade 1 was the largest containing 463/729 total isolates. A majority of Clade 1 isolates were azole-sensitive (292/463) and of the 171 azole-resistant isolates, 80/171 were resistant to more than one azole and only 5 had TR mutations in the *cyp51A* promoter. No Clade 1 isolates had the known mutations associated with resistance to MBC, QoI, or SDHI agricultural fungicides. Clade 2 was the next largest group containing 208/729 isolates. Most Clade 2 isolates were azole-resistant (154/208) with 103 being pan-azole resistant and 128 carrying a TR mutation in the *cyp51A* promoter. Seventy-nine Clade 2 isolates had the known mutations associated with resistance to MBC, QoI, or SDHI agricultural fungicides with 53/79 indicating resistance to more than one agricultural fungicide (multi-fungicide resistant). Interestingly all fungicide-resistant isolates in Clade 2 also contained a TR mutation (79/79). Clade 3 was the smallest (39/729) and contained azole-sensitive (17/39), azole-resistant (20/39), pan-azole-resistant (2/39) and fungicide-resistant isolates (4/39), though no TR or multi-fungicide resistance mutations were present. Among the 729 isolates in our phylogenetic analysis, 19 fell between Clade 1 and Clade 2, suggesting they might be recombinants. Most presumed recombinant isolates were azole resistance (17/19) with 10 being pan-azole resistant. Fourteen isolates among the presumed recombinant isolates carried a TR mutation and were multi-fungicide resistant.

The presence of many TR mutations in Clade 2 and a few in Clade 1 ( C109, C141, C134, and NRZ-2017-214), is similar to findings of previous studies conducted with different isolates (Figure S1) (14, 47). Of the azole-resistant isolates in Clade 1 (Figure S1), roughly 10% had no *cyp51A* mutations and 64% had *cyp51A* mutations that have not been shown to cause azole resistance (A9T, Y46F, V172M, T248N, E255D, K427E, WT) (Table S1). Multi-fungicide resistance was only found in Clade 2 or the presumed recombinant isolates and was mostly associated with isolates with a TR mutation in *cyp51A* (Table 2). There were 5 presumed recombinant isolates with the Y46F/V172M/T248N/E255D/K427E *cyp51A* mutations. Of these, 2 were azole-resistant and 1 was pan-azole-resistant.

**Table 2:** Mutations associated with multi-fungicide resistance in azole-resistant *A. fumigatus*.

| Isolate | Source | Clade | *cyp51A*-azoles | *cytB*-<br>QoI<sup>a</sup> | *benA*-<br>MBC<sup>b</sup> | *sdhB*-<br>SDHI<sup>c</sup> | Reference (DOI) |
| --- | --- | --- | --- | --- | --- | --- | --- |
| Af293 -<br>Bowyer | Clinic, UK | 1 | WT | WT | WT | WT | 10.3390/genes9070363 |
| eAF1438 | Environment,<br>USA | 3 | T248N/E255D/K427E | WT | WT | WT | This study |
| eAF1650 | Environment,<br>USA | 3 | T248N/E255D | G143A | WT | WT | This study |
| AFIS2001 | Clinic, US | 2 | TR34/Y46F/L98H/V172M/T248<br>N/E255D/K427E | G143A | F219Y | WT | 10.1128/mBio.01803-21 |
| Afu<br>1042/09 | Clinic, India | 2 | TR34/Y46F/L98H/V172M/T248<br>N/E255D/K427E | G143A | F219Y | WT | 10.1128/mBio.00536-15 |
| Afu<br>124/E11 | Environment,<br>India | 2 | TR34/Y46F/L98H/V172M/T248<br>N/E255D/K427E | G143A | F219Y | WT | 10.1128/mBio.00536-15 |
| C4 | Environment,<br>Wales | 2 | TR34/Y46F/L98H/V172M/T248<br>N/E255D/K427E | G143A | WT | H270Y | 10.1038/s41564-022-01091-2 |
| C10 | Environment, Wales | 2 | TR34/Y46F/L98H/V172M/T248 N/E255D/K427E | G143A | F219Y | H270R | 10.1038/s41564-022-01091-2 |
| C13 | Environment, Wales | 2 | TR34/Y46F/L98H/V172M/T248 N/E255D/K427E | G143A | F219Y | H270Y | 10.1038/s41564-022-01091-2 |
| C14 | Environment, Wales | 2 | TR34/Y46F/L98H/V172M/T248 N/E255D/K427E | G143A | F219Y | WT | 10.1038/s41564-022-01091-2 |
| C28 | Environment, Wales | 2 | TR34/Y46F/L98H/V172M/T248 N/E255D/K427E | G143A | WT | H270Y | 10.1038/s41564-022-01091-2 |
| C86 | Environment, Ireland | 2 | TR34/Y46F/L98H/V172M/T248 N/E255D/K427E | G143A | WT | H270Y | 10.1038/s41564-022-01091-2 |
| C364 | Environment, Scotland | 2 | TR34/Y46F/L98H/V172M/T248 N/E255D/K427E | G143A | WT | WT | 10.1038/s41564-022-01091-2 |
| eAF1468 | Environment, USA | 2 | TR34/Y46F/L98H/V172M/T248 N/E255D/K427E | G143A | WT | H270Y | This study |
| eAF1472 | Environment, USA | 2 | TR34/Y46F/L98H/V172M/T248 N/E255D/K427E | G143A | F219Y | H270Y | This study |
| eAF1570 | Environment, USA | Recom<sup>d</sup> | TR34/Y46F/L98H/V172M/T248 N/E255D/K427E | G143A | WT | H270Y | This study |
| eAF1668 | Environment, USA | 2 | TR34/Y46F/L98H/V172M/T248 N/E255D/K427E | G143A | F219Y | H270Y | This study |
| eAF1667 | Environment, USA | 2 | TR34/Y46F/L98H/V172M/T248 N/E255D/K427E | WT | F219Y | H270Y | This study |
| eAF1535 | Environment, USA | 2 | TR46/Y46F/Y121F/V172M/T248 N/T298A/K427E | G143A | F219Y | H270Y | This study |
| DI_15-106 | Clinic, USA | 2 | TR46/Y121F/V172M/T248N/T298A/E255D/K427E | G143A | F219Y | H270Y | 10.1128/JCM.02478-15 |
| C155 | Clinic, England | 2 | TR34/Y46F/L98H/V172M/T248 N/E255D/K427E/G448S | G143A | F219Y | WT | 10.1038/s41564-022-01091-2 |
| C89 | Environment, Ireland | 2 | TR46/Y46F/Y121F/V172M/T248 N/E255D/T289A/K427E | G143A | F219Y | WT | 10.1038/s41564-022-01091-2 |
| B11982 | Environment, USA | Recom<sup>d</sup> | TR34/Y46F/L98H/V172M/T248 N/E255D/S297T/K427E/F495I | G143A | F219Y | WT | 10.1128/mBio.01803-21 |
| DI_15-96 | Clinic, USA | 2 | TR46/Y46F/Y121F/V172M/T248 N/E255D/T298A/K427E | G143A | F219Y | WT | 10.1128/JCM.02478-15 |
| eAF222 | Environment, USA | 2 | TR46/Y46F/Y121F/V172M/T248 N/E255D/T298A/K427E | G143A | F219Y | H270Y | 10.1093/g3journal/jkab427 |
| eAF234 | Environment, USA | 2 | TR46/Y46F/Y121F/V172M/T248 N/E255D/T298A/K427E | G143A | F219Y | WT | 10.1093/g3journal/jkab427 |
| eAF513 | Environment, USA | 2 | TR46/Y46F/Y121F/V172M/T248 N/E255D/T298A/K427E | G143A | F219Y | WT | 10.1093/g3journal/jkab427 |
| eAF1458 | Environment, USA | 2 | TR46/Y46F/Y121F/V172M/T248 N/E255D/T298A/K427E | G143A | F219Y | H270Y | This study |
| eAF1685 | Environment, USA | 2 | TR46/Y46F/Y121F/V172M/T248 N/E255D/T298A/K427E | G143A | F219Y | H270R | This study |
| eAF1703 | Environment, USA | 2 | TR46/Y46F/Y121F/V172M/T248 N/E255D/T298A/K427E | G143A | F219Y | WT | This study |
| C93 | Environment, Ireland | 2 | TR46/Y46F/Y121F/V172M/T248 N/E255D/T289A/K427E/G448S | G143A | F219Y | H270Y | 10.1038/s41564-022-01091-2 |

<sup>a</sup> Mutation G143A in *cytB* causes resistance to Qols (49).
<sup>b</sup> Mutation F219Y in *benA* causes resistance to MBCs (50)
<sup>c</sup> Mutation H270Y/R in *sdhB* causes resistance to SDHIs (53)
<sup>d</sup> Recombinant isolate located between Clades 1 and 2 in Figure 2 and Figure S1.

To further investigate population structure, all 729 isolates were included in a principal component analysis (PCA). Isolates congregated in three main clusters supporting a 3-clade structure and presumed recombinant isolates intermixed with Clades 1 and 2, mostly in the area between the two clades (Figure 3).

**Figure 3:**
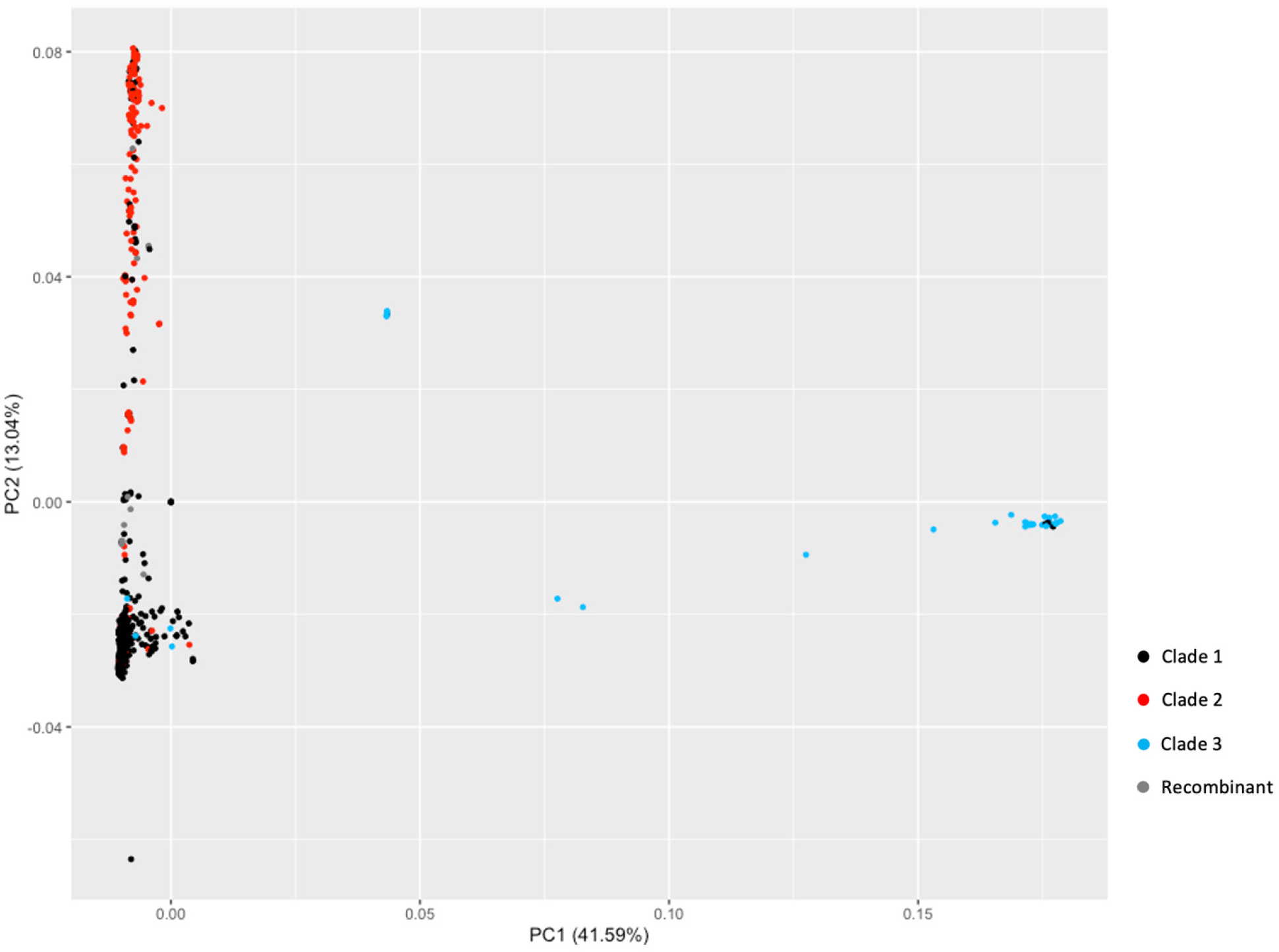
Principal components (PC) 1 and 2 plot mostly separates the three clades. Clade 1 is represented by black dots. Clade 2 is represented by red dots. Clade 3 is represented by blue dots. Presumed recombinant isolates are represented by gray dots. PC 1 represents 41.59% of data along the x-axis. PC 2 represents 13.04% of data on the y-axis.

To investigate recombination between clades, we used our 480 isolate subsample group of *A. fumigatus* U.S. agricultural and worldwide isolates to conduct ADMIXTURE analysis. ADMIXTURE was run with 5 different seeds for K values 2 through 20 and the best log likelihood K values were analyzed (Figure S2). K3 best fit the data and supports 3 clusters that mostly align with the 3 clades (Figure 4). The presumed recombinants showed assignment in both the green and dark purple clusters (Clades 1 and 2).

**Figure 4:**
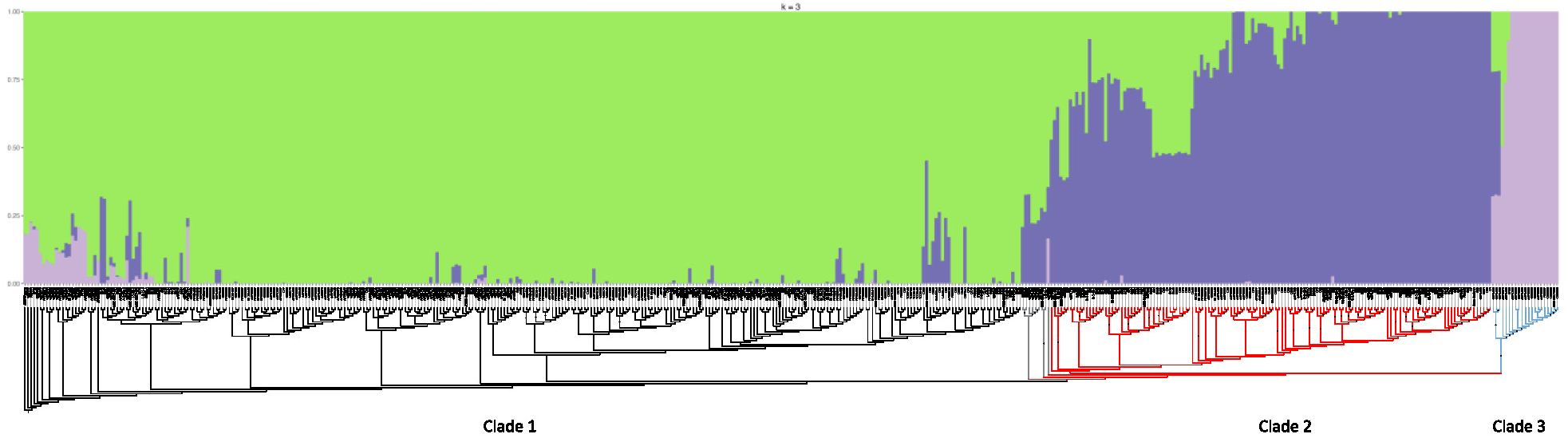
ADMIXTURE analysis supports three populations of *A. fumigatus*. Clade 1 is mostly aligned with the green cluster. Clade 2 is mostly aligned with the dark purple cluster, but several isolates in Clade 2 are also assigned membership to the green cluster. Clade 3 mostly aligns with the light purple cluster but some isolates in this clade show admixture with the green and dark purple clusters. Numbers represent clade number. Y-axis represents ancestry. A rectangular version of the neighbor-joining subsample tree (Figure 2) is on the x-axis. Black branches represent Clade 1. Red branches represent Clade 2. Blue branches represent Clade 3. Gray branches represent presumed recombinant isolates.

Our phylogenetic, ADMIXTURE, and PCA analyses all suggested three populations with the small Clade 3 being the most diverged from the other two. To determine whether Clade 3 might be a cryptic species distinct from *A. fumigatus*, we constructed neighbor-joining gene trees for eight different housekeeping genes (*act1*, *efg1*, *gapdh*, *histH4*, *ITS*, *rpb2*, *sdh1*, and *tef1*) from all 729 isolates (Figure 5 and Figure S3). In no case did Clade 3 form a monophyletic group that diverged from Clades 1 and 2 as would be expected if it were a different species. In 6 of the 8 trees Clade 3 isolates were mostly clustered together, but not monophyletic, and Clade 1 and 2 isolates were interspersed suggesting Clade 3 is rapidly diverging from Clades 1 and 2 (Figure S3). In 2 of the 8 trees, *gapdh* and ITS, isolates from all three clades were interspersed showing less divergence among clades for these specific loci. The absence of monophyly for the gene trees for clade 3 isolates shows current or historically recent gene flow and thus indicates that it is not a distinct species.

**Figure 5:**
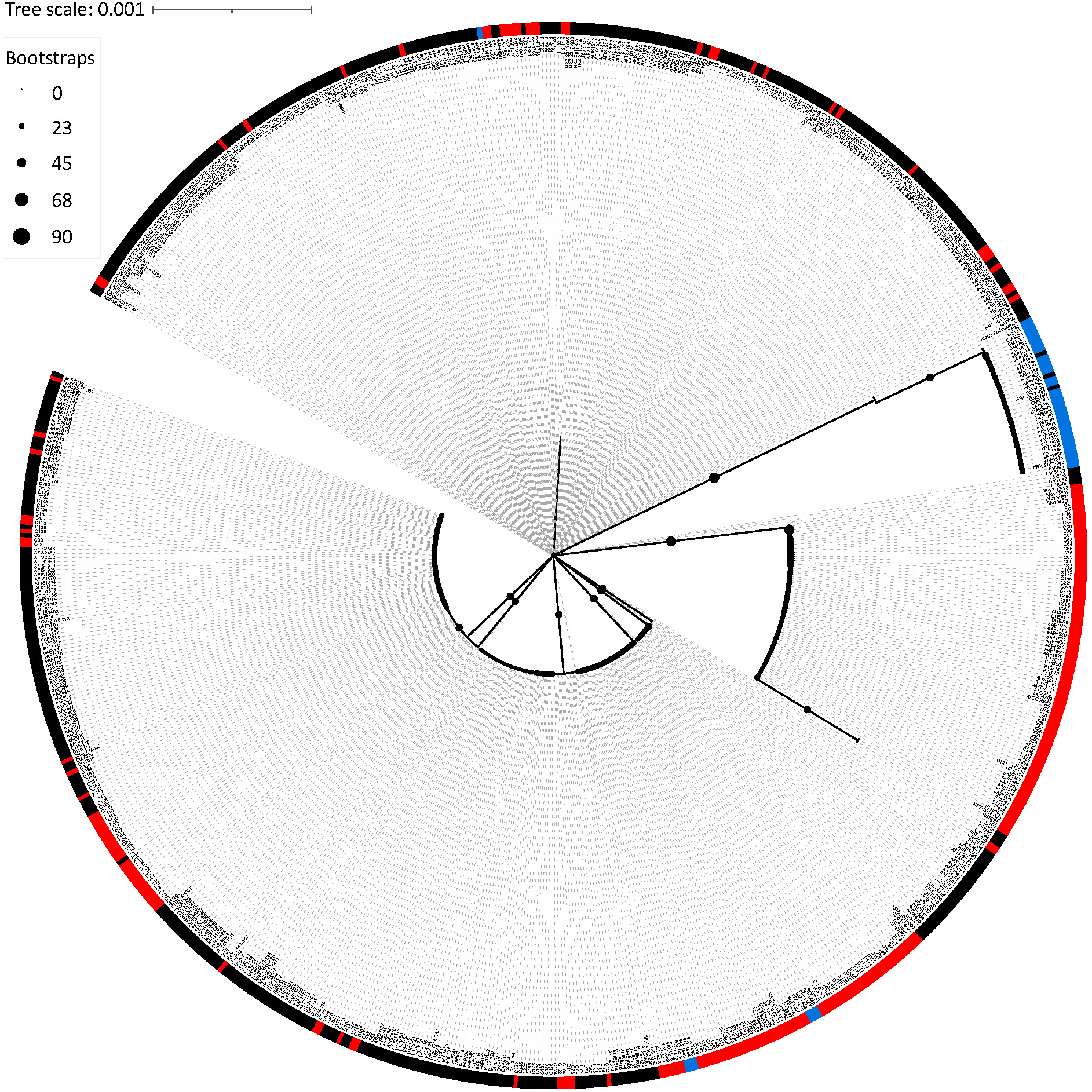
Neighbor-joining tree of housekeeping gene *act1* shows Clade 3 clustering but not monophyletic. A gene tree of *act1* from 729 isolates was constructed. Circle size on branches are a proxy for bootstraps. Black squares represent isolates from Clade 1. Red squares represent isolates from Clade 2. Blue squares represent isolates from Clade 3.

### Recombinant strains of *A. fumigatus*

The 19 isolates that appeared to be recombinants with Clade 1 and Clade 2 ancestry showed a variety of azole resistance phenotypes and other fungicide resistance genotypes. To determine if Clade 1 or Clade 2 contributed the Chromosome 4 *cyp51A* allele in the azole resistant recombinant strains, we used the faChrompaint.pl script to paint 6 isolates’ chromosomes *in silico* (39). We found all 6 recombinant isolates inherited the *cyp51A* allele conferring azole resistance from Clade 2 ancestors (Figure 6).

**Figure 6:**
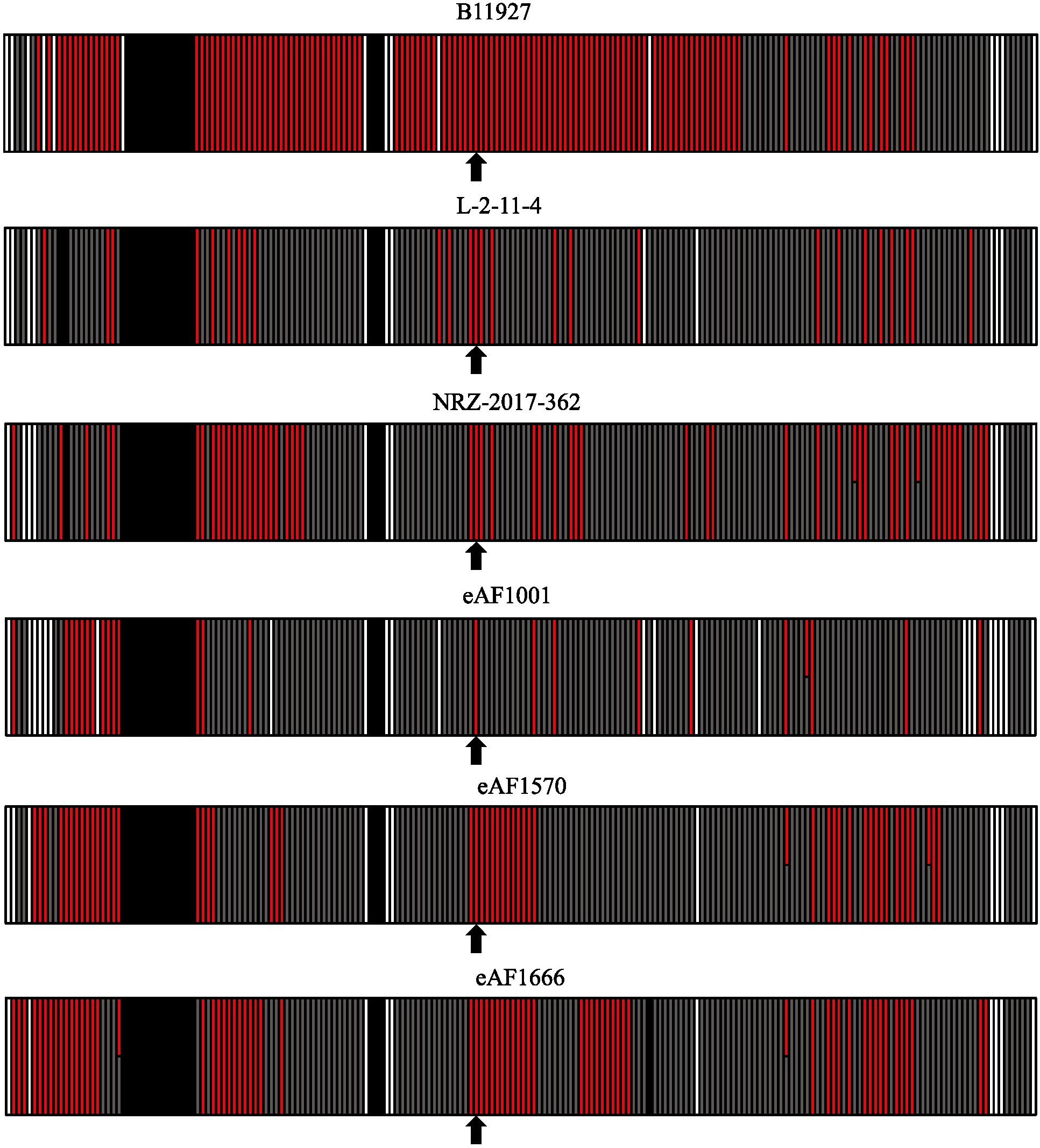
The *cyp51A* locus came from Clade 2 in azole-resistant admixed strains of *A. fumigatus*. Chromosome 4 is painted gray for Clade 1 origin, red for Clade 2 origin, black for low quality sequencing, and white for areas too far diverged from Clade 1 and Clade 2. Arrows represent the window the *cyp51A* locus. Window size is 5kb.

## Discussion

### Azole-resistant *A. fumigatus* is widespread in the US

Before the current study *A. fumigatus* isolates from clinical sources had been reported in 37 states and from environmental sources in 2 states (Georgia and Florida) (11, 20, 54) (Figure 1). We surveyed agricultural sources in 7 additional states including 4 on the eastern U.S. coast and 3 on the western U.S. coast. We combined our data with publicly available data to get an overview of *A. fumigatus* reported from U.S. clinical and environmental sources. Sampling intensity has varied widely so we could not determine the true prevalence of azole-resistant *A. fumigatus* in each state or associated with different crops. However, it is clear that azole-resistant *A. fumigatus* is widespread in the U.S. We found it in agricultural environments on both coasts where a variety of crop types have been cultivated. Interestingly, we found high levels of TR-based azole resistance and multi-fungicide resistance in isolates from tulip farms on the west coast that also grow hemp (Supplementary table 1). Most tulips in the U.S. are grown from bulbs imported from the Netherlands, where TR-based azole resistance was first reported and is thought to have originated (55, 56).

### Our data support 3 clades of *Aspergillus fumigatus* worldwide

Previous work from multiple labs showed that worldwide *A. fumigatus* populations were not structured by geography (41, 57, 58). Most recent work using clinical and environmental isolates suggested that *A. fumigatus* populations fell into two major clades with most TR-based pan-azole-resistance in Clade A (11–13). A notable exception is pan-genome analysis which showed 7 genetic clusters of *A. fumigatus*, though this analysis also found close clustering of TR-based azole resistance and no strong geographic structure (47). A recent pan-genome analysis with increased sampling identified three clades: a large group with few azole-resistant isolates called Clade 1 by the authors (equivalent to Clade B in earlier work), a somewhat smaller group of isolates that included most isolates with TR-based pan-azole resistance called Clade 2 by the authors (equivalent to Clade A in earlier work) and a third clade with the fewest number of isolates and no azole resistance called Clade 3 by the authors (16). In the current study, our phylogenetic, PCA and ADMIXTURE analyses of 729 U.S. and worldwide isolates support three clades: Clade 1 with 463 isolates (equivalent to Clade B in previous work), Clade 2 with 208 isolates (equivalent to Clade A in previous work) and the small Clade 3 with 39 isolates (Figures 2, 3, and 4). To ensure that Clade 3 was not a cryptic species we performed phylogenetic analyses with 8 housekeeping genes. In all cases the tree topology was inconsistent with monophyly of Clade 3, which would be expected for a cryptic species. The clustering of Clade 3 isolates among trees supported the presence of a third clade of *A. fumigatus*, but not a cryptic species (Figure 5 and Figure S3).

### TR-based pan azole resistance and multi-fungicide resistance are strongly associated

We found that all but 5 isolates that contained a TR mutation in the *cyp51A* promoter were in Clade 2 and found no TR-based azole resistance in Clade 3, similar to previous reports which included relatively few U.S. isolates (16). Interestingly, none of the 83 isolates resistant to the agricultural fungicides MBC, QoI, or SDH were in Clade 1. A total of 79 fungicide-resistant isolates were in Clade 2 and all 79 also carried either the TR_34_ or the TR_46_ azole-resistance alleles of *cyp51A* (Figure S1, Table S2). This pattern suggests that the genetic background of Clade 2 isolates allows more rapid adaptation to environmental stress.

Though we found no azole resistance in Clade 3, we did find the G143A *cytB* mutation that causes resistance to the agricultural QoI fungicides in 4 of the 39 isolates (Table 2, Table S2). Based on phylogenetic analysis of whole genome sequence (Figure 2 and Figure S1) and individual housekeeping genes (Figure 5 and Figure S3), Clade 3 appears to be recently diverged from Clades 1 and 2. Because the number of Clade 3 isolates is relatively small, it is possible that azole-resistant isolates from this clade will be discovered in the future.

### U.S. populations are recombining

Previous work from others found evidence of very high recombination rates in *A. fumigatus* with recombination frequently occurring within clades (16, 59). Our isolates also showed recombination and admixture, with a notable group falling between Clades 1 and 2 (Figure 2, Figure S1). Interestingly 17 of the 19 recombinants from this group were environmental isolates from the U.S. with the remaining 2 being from Germany, one from soil and one from a patient (Figure S1, Table S1). Ten of the isolates were pan-azole resistant. PCA showed individuals in this group as overlapping with Clades 1 and 2, but mostly in the area between Clades 1 and 2 (Figure 3). ADMIXTURE analysis was also consistent with three recombining populations (Figure 4). Similar to work from others (60), higher K values increased sub-structuring of the populations (Figure S2). Chromosome painting for 6 of the recombinant isolates showed that the pan-azole resistance-allele derived from Clade 2 (Figure 6).

## Conclusion

Our study expands environmental sampling of *A. fumigatus* in the U.S. We show that azole-resistant and multi-fungicide-resistant isolates are found in agricultural settings in both the east and west coast regions of the country in areas where a variety of crops have been cultivated. We show support for three clades of *A. fumigatus*, consistent with a recent study that included many fewer U.S. isolates (Figure 3) (16). We found the small Clade 3 has no TR-based azole resistance but does contain isolates with QoI fungicide resistance. Interestingly, the remaining 79 isolates that were resistant to agricultural fungicides MBC, QoI, or SDHI were in Clade 2 and all also carried a TR allele of *cyp51A* associated with pan-azole resistance. U.S. isolates also showed high levels of recombination raising the worrying prospect that resistance to clinical azoles will continue to spread.

## Acknowledgements

This work was supported by the Centers for Disease Control and Prevention (CDC; contract 0HCVLD13-2018-27470 to M.M. and M.T.B.) and United States Department of Agriculture, National Institute of Food and Agriculture (USDA NIFA AFRI grant 2019-67017-29113 to M.T.B. and M.M.). B.N.C-S. was also supported by the National Science Foundation under Grant No. DGE-1545433.

## References

1. World Health Organization. 2022. WHO fungal priority pathogens list to guide research, development and public health action. https://www.who.int/publications/i/item/9789240060241

2. Bongomin F, Gago S, Oladele RO, Denning DW. 2017. Global and Multi-National Prevalence of Fungal Diseases-Estimate Precision. J Fungi (Basel) 3.

3. Patterson TF, Thompson GR, 3rd, Denning DW, Fishman JA, Hadley S, Herbrecht R, Kontoyiannis DP, Marr KA, Morrison VA, Nguyen MH, Segal BH, Steinbach WJ, Stevens DA, Walsh TJ, Wingard JR, Young JA, Bennett JE. 2016. Practice Guidelines for the Diagnosis and Management of Aspergillosis: 2016 Update by the Infectious Diseases Society of America. Clin Infect Dis 63:e1–e60.

4. Klittich C, Jr., Green FR, 3rd, Ruiz JM, Weglarz T, Blakeslee BA. 2008. Assessment of fungicide systemicity in wheat using LC-MS/MS. Pest Manag Sci 64:1267–77.

5. Celia-Sanchez BN, Mangum B, Brewer M, Momany M. 2022. Analysis of Cyp51 protein sequences shows 4 major Cyp51 gene family groups across fungi. G3 (Bethesda) 12.

6. Marichal P, Gorrens J, Laurijssens L, Vermuyten K, Van Hove C, Le Jeune L, Verhasselt P, Sanglard D, Borgers M, Ramaekers FC, Odds F, Vanden Bossche H. 1999. Accumulation of 3-ketosteroids induced by itraconazole in azole-resistant clinical Candida albicans isolates. Antimicrob Agents Chemother 43:2663–70.

7. Alvarez-Moreno C, Lavergne RA, Hagen F, Morio F, Meis JF, Le Pape P. 2017. Azole-resistant Aspergillus fumigatus harboring TR(34)/L98H, TR(46)/Y121F/T289A and TR(53) mutations related to flower fields in Colombia. Sci Rep 7:45631.

8. Vermeulen E, Maertens J, Schoemans H, Lagrou K. 2012. Azole-resistant Aspergillus fumigatus due to TR46/Y121F/T289A mutation emerging in Belgium, July 2012. Euro Surveill 17.

9. Mellado E, Garcia-Effron G, Alcazar-Fuoli L, Melchers WJ, Verweij PE, Cuenca-Estrella M, Rodriguez-Tudela JL. 2007. A new Aspergillus fumigatus resistance mechanism conferring in vitro cross-resistance to azole antifungals involves a combination of cyp51A alterations. Antimicrob Agents Chemother 51:1897–904.

10. Burks C, Darby A, Gomez Londono L, Momany M, Brewer MT. 2021. Azole-resistant Aspergillus fumigatus in the environment: Identifying key reservoirs and hotspots of antifungal resistance. PLoS Pathog 17:e1009711.

11. Kang SE, Sumabat LG, Melie T, Mangum B, Momany M, Brewer MT. 2022. Evidence for the agricultural origin of resistance to multiple antimicrobials in Aspergillus fumigatus, a fungal pathogen of humans. G3 (Bethesda) 12.

12. Sewell TR, Zhu J, Rhodes J, Hagen F, Meis JF, Fisher MC, Jombart T. 2019. Nonrandom Distribution of Azole Resistance across the Global Population of Aspergillus fumigatus. mBio 10.

13. Etienne KA, Berkow EL, Gade L, Nunnally N, Lockhart SR, Beer K, Jordan IK, Rishishwar L, Litvintseva AP. 2021. Genomic Diversity of Azole-Resistant Aspergillus fumigatus in the United States. Mbio 12.

14. Rhodes J, Abdolrasouli A, Dunne K, Sewell TR, Zhang Y, Ballard E, Brackin AP, van Rhijn N, Chown H, Tsitsopoulou A, Posso RB, Chotirmall SH, McElvaney NG, Murphy PG, Talento AF, Renwick J, Dyer PS, Szekely A, Bowyer P, Bromley MJ, Johnson EM, Lewis White P, Warris A, Barton RC, Schelenz S, Rogers TR, Armstrong-James D, Fisher MC. 2022. Population genomics confirms acquisition of drug-resistant Aspergillus fumigatus infection by humans from the environment. Nat Microbiol 7:663–674.

15. Pinto E, Monteiro C, Maia M, Faria MA, Lopes V, Lameiras C, Pinheiro D. 2018. Aspergillus Species and Antifungals Susceptibility in Clinical Setting in the North of Portugal: Cryptic Species and Emerging Azoles Resistance in A. fumigatus. Front Microbiol 9:1656.

16. Lofgren LA, Ross BS, Cramer RA, Stajich JE. 2022. The pan-genome of Aspergillus fumigatus provides a high-resolution view of its population structure revealing high levels of lineage-specific diversity driven by recombination. PLoS Biol 20:e3001890.

17. Fan Y, Wang Y, Korfanty GA, Archer M, Xu J. 2021. Genome-Wide Association Analysis for Triazole Resistance in Aspergillus fumigatus. Pathogens 10.

18. Snelders E, Huis In ’t Veld RA, Rijs AJ, Kema GH, Melchers WJ, Verweij PE. 2009. Possible environmental origin of resistance of Aspergillus fumigatus to medical triazoles. Appl Environ Microbiol 75:4053–7.

19. Verweij PE, Howard SJ, Melchers WJ, Denning DW. 2009. Azole-resistance in Aspergillus: proposed nomenclature and breakpoints. Drug Resist Updat 12:141–7.

20. Hurst SF, Berkow EL, Stevenson KL, Litvintseva AP, Lockhart SR. 2017. Isolation of azole-resistant Aspergillus fumigatus from the environment in the south-eastern USA. J Antimicrob Chemother 72:2443–2446.

21. Arendrup MC, Friberg N, Mares M, Kahlmeter G, Meletiadis J, Guinea J, Subcommittee on Antifungal Susceptibility Testing of the EECfAST. 2020. How to interpret MICs of antifungal compounds according to the revised clinical breakpoints v. 10.0 European committee on antimicrobial susceptibility testing (EUCAST). Clin Microbiol Infect 26:1464–1472.

22. Camps SMT, Dutilh BE, Arendrup MC, Rijs AJMM, Snelders E, Huynen MA, Verweij PE, Melchers WJG. 2012. Discovery of a hapE Mutation That Causes Azole Resistance in Aspergillus fumigatus through Whole Genome Sequencing and Sexual Crossing. Plos One 7.

23. Pitkin JW, Panaccione DG, Walton JD. 1996. A putative cyclic peptide efflux pump encoded by the TOXA gene of the plant-pathogenic fungus Cochliobolus carbonum. Microbiology-Uk 142:1557–1565.

24. Bankevich A, Nurk S, Antipov D, Gurevich AA, Dvorkin M, Kulikov AS, Lesin VM, Nikolenko SI, Pham S, Prjibelski AD, Pyshkin AV, Sirotkin AV, Vyahhi N, Tesler G, Alekseyev MA, Pevzner PA. 2012. SPAdes: a new genome assembly algorithm and its applications to single-cell sequencing. J Comput Biol 19:455–77.

25. Altschul SF, Gish W, Miller W, Myers EW, Lipman DJ. 1990. Basic local alignment search tool. J Mol Biol 215:403–10.

26. Quinlan AR, Hall IM. 2010. BEDTools: a flexible suite of utilities for comparing genomic features. Bioinformatics 26:841–2.

27. Li H, Durbin R. 2009. Fast and accurate short read alignment with Burrows-Wheeler transform. Bioinformatics 25:1754–60.

28. Li H, Handsaker B, Wysoker A, Fennell T, Ruan J, Homer N, Marth G, Abecasis G, Durbin R, Genome Project Data Processing S. 2009. The Sequence Alignment/Map format and SAMtools. Bioinformatics 25:2078–9.

29. Li H. 2011. A statistical framework for SNP calling, mutation discovery, association mapping and population genetical parameter estimation from sequencing data. Bioinformatics 27:2987–93.

30. Stecher G, Tamura K, Kumar S. 2020. Molecular Evolutionary Genetics Analysis (MEGA) for macOS. Mol Biol Evol 37:1237–1239.

31. Letunic I, Bork P. 2019. Interactive Tree Of Life (iTOL) v4: recent updates and new developments. Nucleic Acids Res 47:W256–W259.

32. Kimura M. 1980. A simple method for estimating evolutionary rates of base substitutions through comparative studies of nucleotide sequences. J Mol Evol 16:111–20.

33. Price AL, Patterson NJ, Plenge RM, Weinblatt ME, Shadick NA, Reich D. 2006. Principal components analysis corrects for stratification in genome-wide association studies. Nat Genet 38:904–9.

34. Team RC. 2021. A Language and Environment for Statistical Computing R Foundation for Statistical Computing. https://www.R-project.org/. Accessed

35. Wickham H. 2016. ffplot2: Elegant Graphics for Data Analysis. Springer-Verlag New York.

36. Purcell S, Neale B, Todd-Brown K, Thomas L, Ferreira MA, Bender D, Maller J, Sklar P, de Bakker PI, Daly MJ, Sham PC. 2007. PLINK: a tool set for whole-genome association and population-based linkage analyses. Am J Hum Genet 81:559–75.

37. Alexander DH, Novembre J, Lange K. 2009. Fast model-based estimation of ancestry in unrelated individuals. Genome Res 19:1655–64.

38. Kopelman NM, Mayzel J, Jakobsson M, Rosenberg NA, Mayrose I. 2015. Clumpak: a program for identifying clustering modes and packaging population structure inferences across K. Mol Ecol Resour 15:1179–91.

39. Bensasson D, Dicks J, Ludwig JM, Bond CJ, Elliston A, Roberts IN, James SA. 2019. Diverse Lineages of Candida albicans Live on Old Oaks. Genetics 211:277–288.

40. Howard SJ, Pasqualotto AC, Denning DW. 2010. Azole resistance in allergic bronchopulmonary aspergillosis and Aspergillus bronchitis. Clin Microbiol Infect 16:683–8.

41. Abdolrasouli A, Rhodes J, Beale MA, Hagen F, Rogers TR, Chowdhary A, Meis JF, Armstrong-James D, Fisher MC. 2015. Genomic Context of Azole Resistance Mutations in Aspergillus fumigatus Determined Using Whole-Genome Sequencing. mBio 6:e00536.

42. Wiederhold NP, Gil VG, Gutierrez F, Lindner JR, Albataineh MT, McCarthy DI, Sanders C, Fan H, Fothergill AW, Sutton DA. 2016. First Detection of TR34 L98H and TR46 Y121F T289A Cyp51 Mutations in Aspergillus fumigatus Isolates in the United States. J Clin Microbiol 54:168–71.

43. Ballard E, Melchers WJG, Zoll J, Brown AJP, Verweij PE, Warris A. 2018. In-host microevolution of Aspergillus fumigatus: A phenotypic and genotypic analysis. Fungal Genet Biol 113:1–13.

44. Garcia-Rubio R, Monzon S, Alcazar-Fuoli L, Cuesta I, Mellado E. 2018. Genome-Wide Comparative Analysis of Aspergillus fumigatus Strains: The Reference Genome as a Matter of Concern. Genes (Basel) 9.

45. Rybak JM, Ge W, Wiederhold NP, Parker JE, Kelly SL, Rogers PD, Fortwendel JR. 2019. Mutations in hmg1, Challenging the Paradigm of Clinical Triazole Resistance in Aspergillus fumigatus. mBio 10.

46. Dos Santos RAC, Steenwyk JL, Rivero-Menendez O, Mead ME, Silva LP, Bastos RW, Alastruey-Izquierdo A, Goldman GH, Rokas A. 2020. Genomic and Phenotypic Heterogeneity of Clinical Isolates of the Human Pathogens Aspergillus fumigatus, Aspergillus lentulus, and Aspergillus fumigatiaffinis. Front Genet 11:459.

47. Barber AE, Sae-Ong T, Kang K, Seelbinder B, Li J, Walther G, Panagiotou G, Kurzai O. 2021. Aspergillus fumigatus pan-genome analysis identifies genetic variants associated with human infection. Nat Microbiol 6:1526–1536.

48. Steenwyk JL, Mead ME, de Castro PA, Valero C, Damasio A, Dos Santos RAC, Labella AL, Li Y, Knowles SL, Raja HA, Oberlies NH, Zhou X, Cornely OA, Fuchs F, Koehler P, Goldman GH, Rokas A. 2021. Genomic and Phenotypic Analysis of COVID-19-Associated Pulmonary Aspergillosis Isolates of Aspergillus fumigatus. Microbiol Spectr 9:e0001021.

49. Gisi U. 2014. Assessment of selection and resistance risk for demethylation inhibitor fungicides in Aspergillus fumigatus in agriculture and medicine: a critical review. Pest Manag Sci 70:352–64.

50. Hawkins NJ, Fraaije BA. 2016. Predicting Resistance by Mutagenesis: Lessons from 45 Years of MBC Resistance. Front Microbiol 7:1814.

51. Fraaije B, Atkins S, Hanley S, Macdonald A, Lucas J. 2020. The Multi-Fungicide Resistance Status of Aspergillus fumigatus Populations in Arable Soils and the Wider European Environment. Front Microbiol 11:599233.

52. Gonzalez-Jimenez I, Garcia-Rubio R, Monzon S, Lucio J, Cuesta I, Mellado E. 2021. Multiresistance to Nonazole Fungicides in Aspergillus fumigatus TR(34)/L98H Azole-Resistant Isolates. Antimicrob Agents Chemother 65:e0064221.

53. Fraaije BA, Bayon C, Atkins S, Cools HJ, Lucas JA, Fraaije MW. 2012. Risk assessment studies on succinate dehydrogenase inhibitors, the new weapons in the battle to control Septoria leaf blotch in wheat. Mol Plant Pathol 13:263–75.

54. Etienne KA, Berkow EL, Gade L, Nunnally N, Lockhart SR, Beer K, Jordan IK, Rishishwar L, Litvintseva AP. 2021. Genomic Diversity of Azole-Resistant Aspergillus fumigatus in the United States. mBio 12:e0180321.

55. Hagiwara D. 2020. Isolation of azole-resistant Aspergillus fumigatus from imported plant bulbs in Japan and the effect of fungicide treatment. J Pestic Sci 45:147–150.

56. Dunne K, Hagen F, Pomeroy N, Meis JF, Rogers TR. 2017. Intercountry Transfer of Triazole-Resistant Aspergillus fumigatus on Plant Bulbs. Clin Infect Dis 65:147–149.

57. Klaassen CH, Gibbons JG, Fedorova ND, Meis JF, Rokas A. 2012. Evidence for genetic differentiation and variable recombination rates among Dutch populations of the opportunistic human pathogen Aspergillus fumigatus. Mol Ecol 21:57–70.

58. Ashu EE, Hagen F, Chowdhary A, Meis JF, Xu J. 2017. Global Population Genetic Analysis of Aspergillus fumigatus. mSphere 2.

59. Auxier B, Becker F, Nijland R, Debets A, van den Heuvel J, Snelders E. 2022. Meiosis in the human pathogen Aspergillus fumigatus has the highest known number of crossovers. bioRxiv.

60. Wei X, Chen P, Gao R, Li Y, Zhang A, Liu F, Lu L. 2017. Screening and Characterization of a Non-cyp51A Mutation in an Aspergillus fumigatus cox10 Strain Conferring Azole Resistance. Antimicrob Agents Chemother 61.

