## supplemental material for "Pan-azole- and multi-fungicide-resistant *Aspergillus fumigatus* is widespread in the United States"

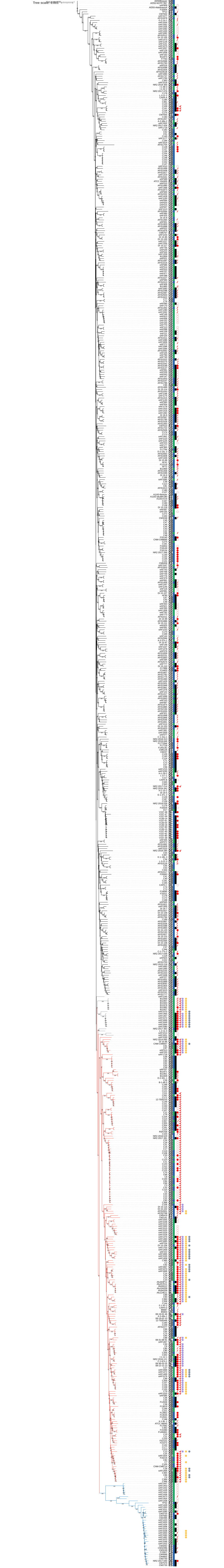
Tree scale: 0.001

**Supplemental Figure 1: Neighbor-joining tree of 729 environmental and clinical isolates of *Aspergillus fumigatus*.** Whole genome sequences from agricultural sites in the east and west coast regions of the United States (eAF1XXX) were analyzed with publicly available data. Af293 was used as the reference genome and root of the tree. Bootstrap values are indicated on each branch. Clade 1 branches are black; Clade 2 branches are red; Clade 3 branches are blue; and branches for strains recombining between Clades 1 and 2 are gray. Country of origin is listed next to each isolate according to their two-letter designation (CA, Canada; DE, Germany; ES, Spain; GB, Great Britain; IN, India; NL, Netherlands; US, United States). Green and blue bars indicate environmental and clinical isolates, respectively. Black and white bars represent mating type MAT1-1 and MAT1-2, respectively. Open red circles indicate azole resistant isolates. Solid red circles indicate pan-azole resistant isolates (i.e., isolates resistant to at least two different clinical azoles based on MIC testing). Red slash marks represent isolates without MIC test data for either itraconazole, voriconazole, or posaconazole. Red check marks indicate isolates with TR mutations. Yellow circles indicate the cytB G143A mutation conferring resistance to QoI fungicides (Gisi 2014). Purple circles indicate the benA F219Y mutation conferring resistance to MBC fungicides (Hawkins and Fraaije 2016). Gray circles indicate the sdhB H270Y/R mutations conferring resistance to SDHI fungicides (Fraaije et al. 2012).

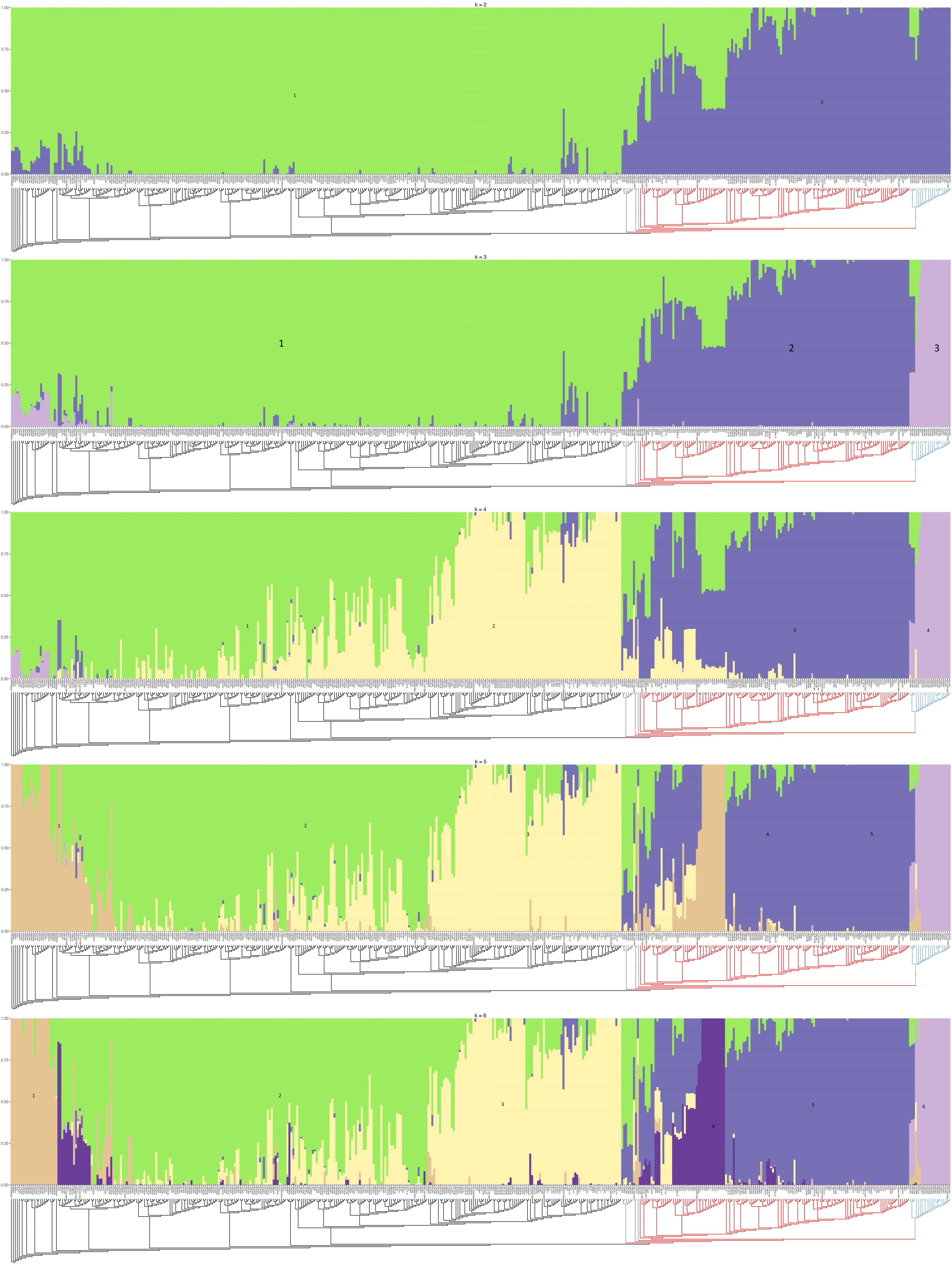
k = 2
1.00
0.75
0.50
0.25
0.00
1
2
k = 3
1
2
3
k = 4
1
2
3
4
k = 5
1
2
3
4
5
k = 6
1
2
3
4
5
6

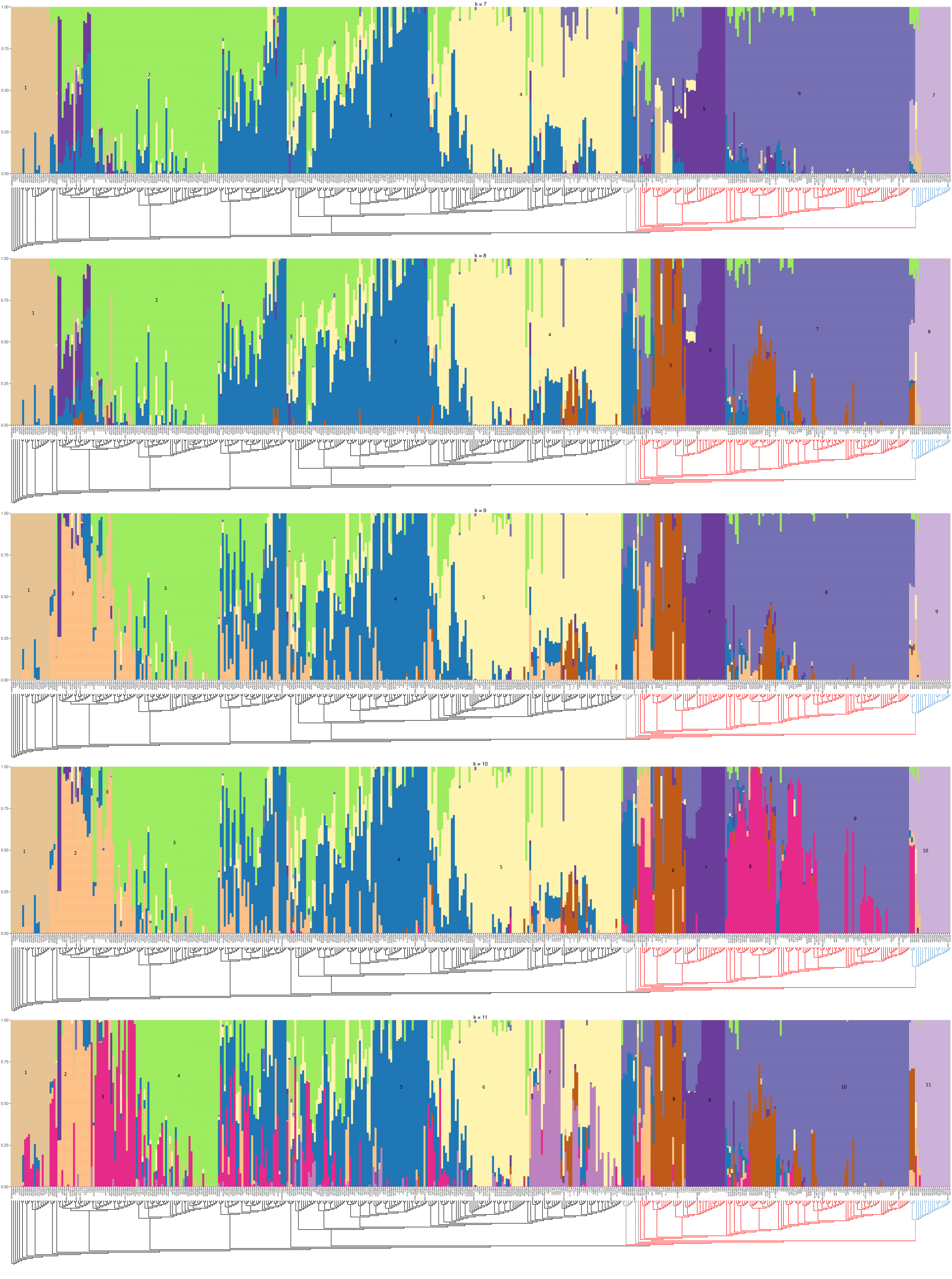

k = 7
1.00
0.75
0.50
0.25
0.00
1
2
3
4
5
6
7
k = 8
1
2
3
4
5
6
7
8
k = 9
1
2
3
4
5
6
7
8
9
k = 10
1
2
3
4
5
6
7
8
9
10
k = 11
1
2
3
4
5
6
7
8
9
10
11

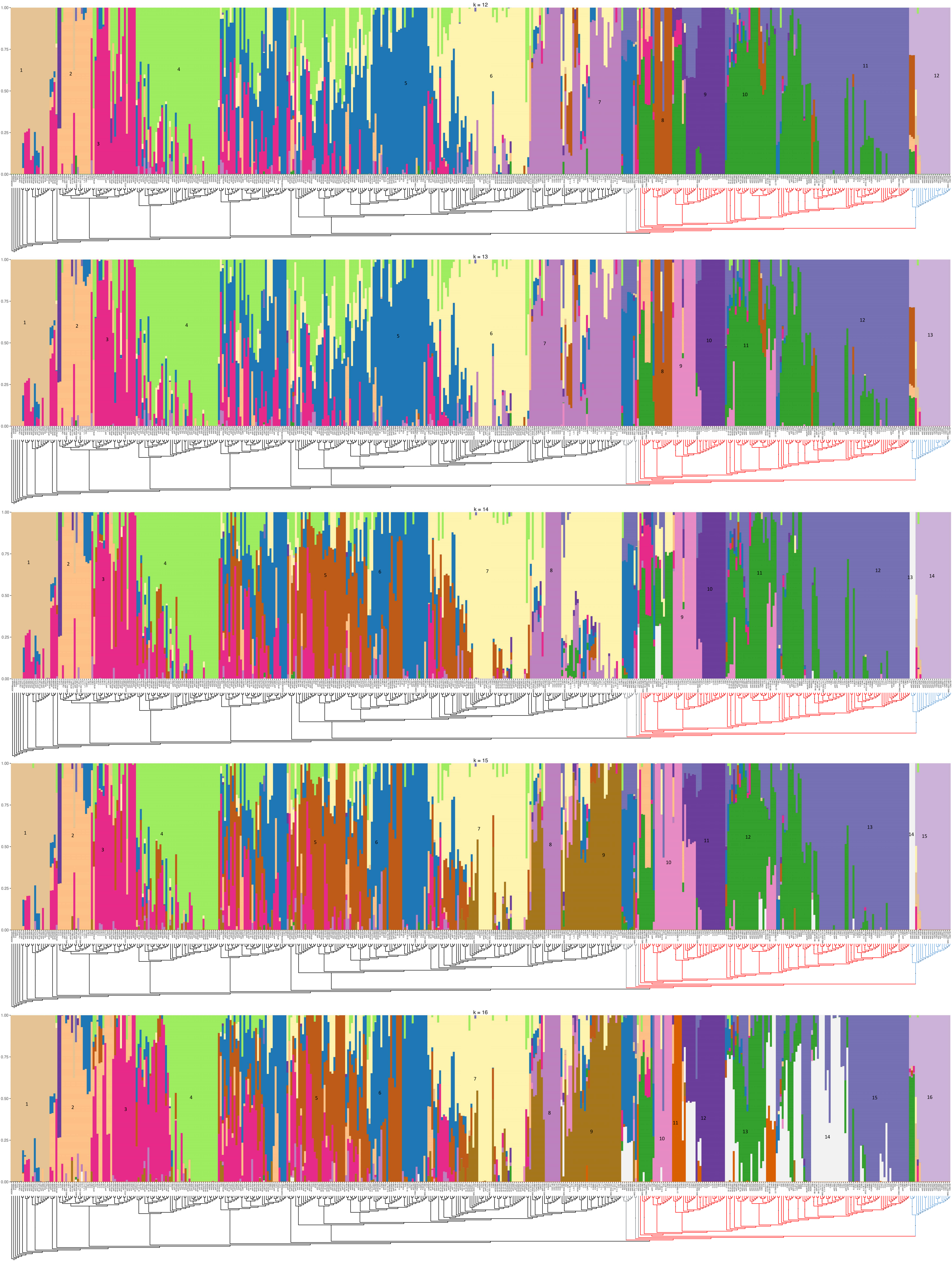

k = 12
k = 13
k = 14
k = 15
k = 16

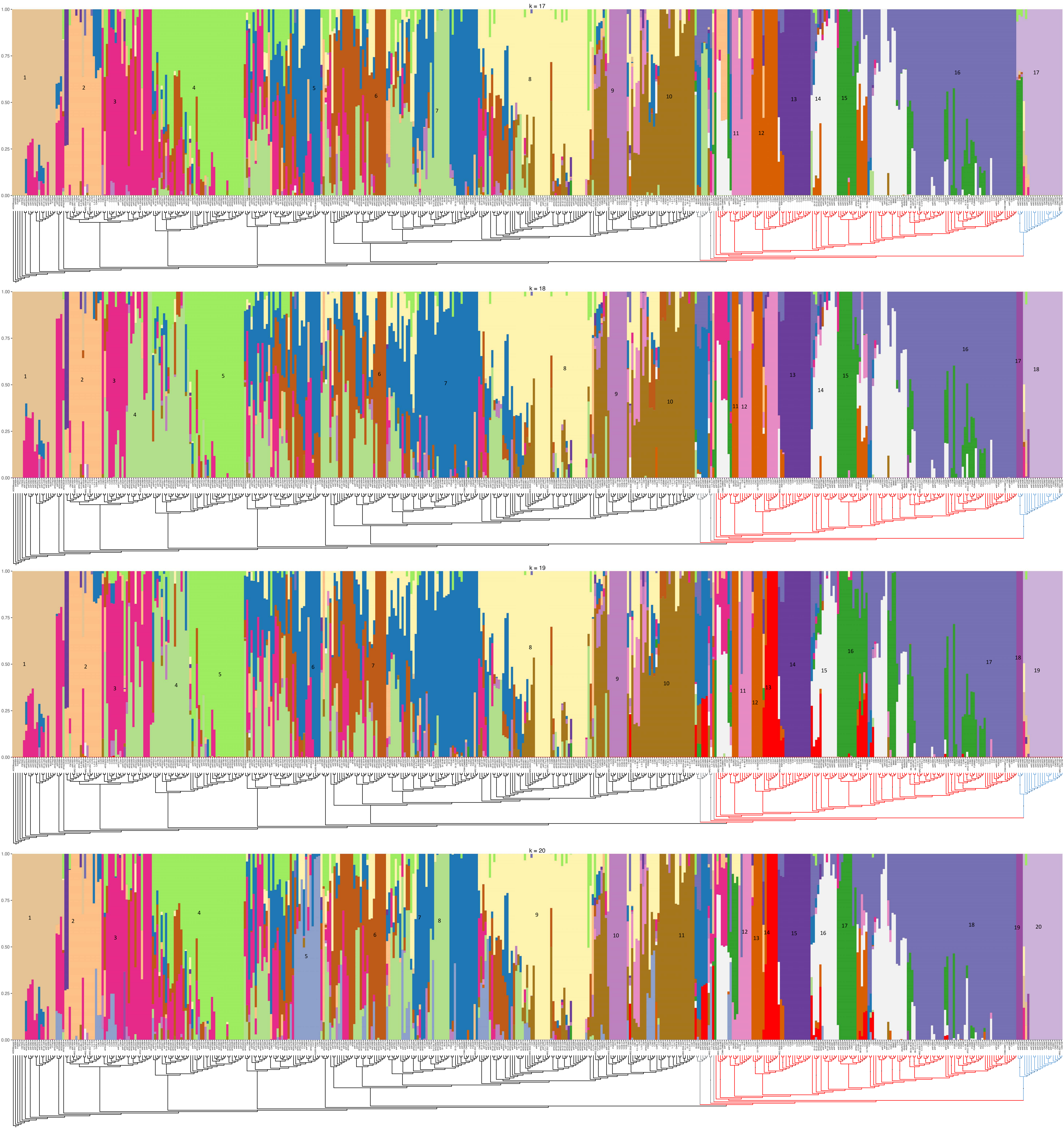
k = 17
k = 18
k = 19
k = 20
1.00
0.75
0.50
0.25
0.00

**Supplemental Figure 2: ADMIXTURE analysis of K2-20.** Numbers within plots represent populations. Y-axis represents ancestry. A rectangular version of the neighbor-joining subsample tree (Figure 2) is on the x-axis. Black branches represent Clade 1. Red branches represent Clade 2. Blue branches represent Clade 3. Gray branches represent presumed recombinant isolates.

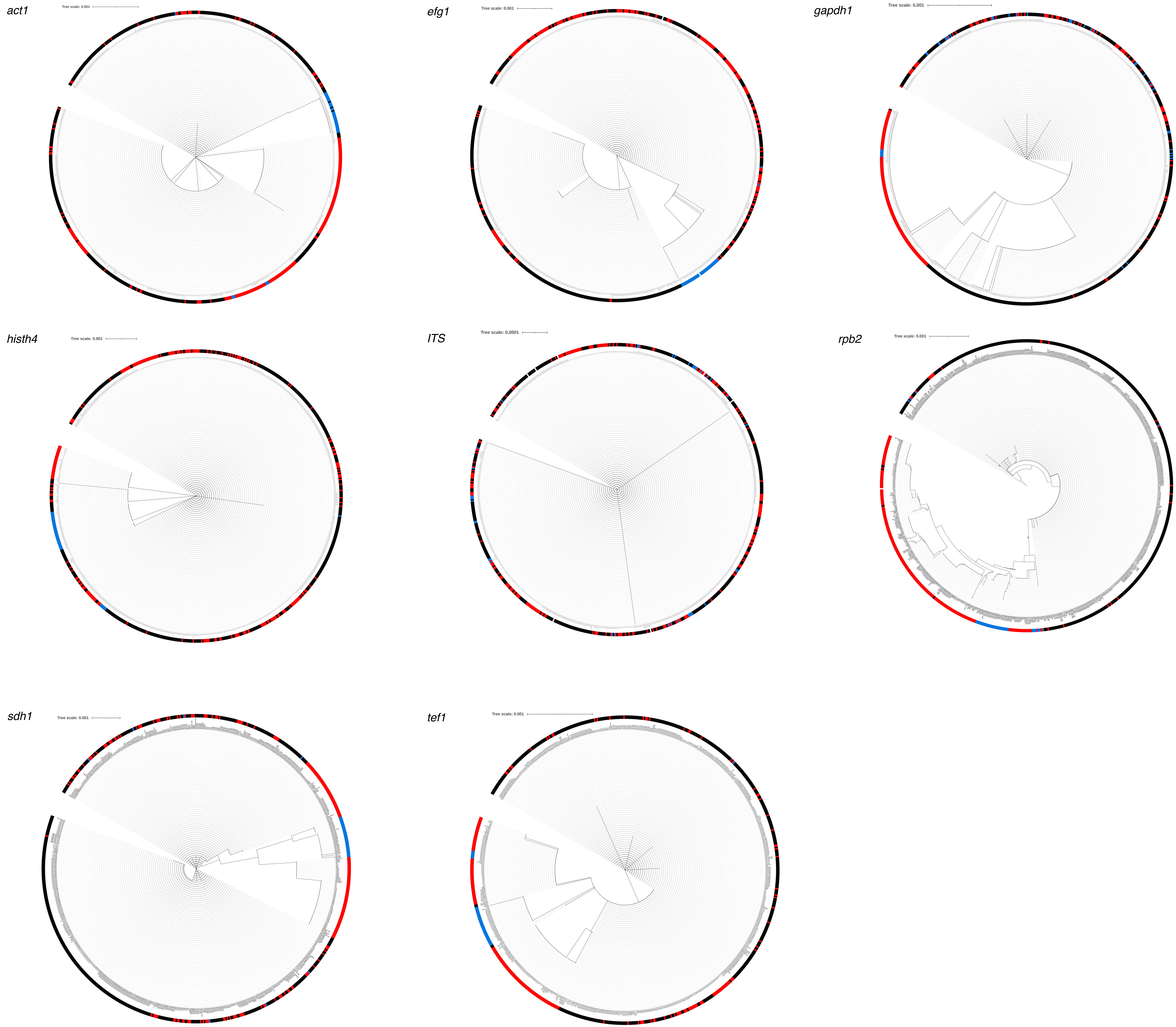
act1
Tree scale: 0.001
efg1
Tree scale: 0.001
gapdh1
Tree scale: 0.001
histh4
Tree scale: 0.001
ITS
Tree scale: 0.0001
rpb2
Tree scale: 0.001
sdh1
Tree scale: 0.001
tef1
Tree scale: 0.001

**Supplemental Figure 3: Neighbor-joining trees of housekeeping genes and ITS region.** Gene trees with 729 isolates were constructed for *act1, efg1, gapdh1, histh4,* the ITS region*, rpb2, sdh1,* and *tef1.* Circle size on branches is a proxy for bootstrap values. Black squares represent isolates from Clade 1. Red squares represent isolates from Clade 2. Blue squares represent isolates from Clade 3.
